# The transformer as a candidate canonical cortical microcircuit

**DOI:** 10.64898/2026.11.01.763457

**Authors:** Yash Shah, Daniel L. K. Yamins

## Abstract

Neuroscience hypothesizes a canonical microcircuit subserving cortical computation across brain systems. In artificial intelligence, neural networks have become a universal computational substrate. Here, we leverage newly available large-scale data and mapping techniques to compare neural network motifs to the brain. We embedded a spectrum of candidate microcircuit architectures within a common spatial macrocircuit, measuring alignment with an extensive battery of cortical signatures. We find that one specific circuit type—the transformer—uniquely captures fine-grained unit-level circuit details, population-level functional maps and neuronal activity patterns, and systems-level wiring efficiency and behavioral demands. The key ingredient enabling the transformer’s success is a learned multiplicative component making relational information available without extensive long-range wiring. These results suggest striking convergence between computational motifs in natural and artificial intelligence.

## Introduction

The cortex supports an extraordinary diversity of behaviors despite sharing remarkably conserved local cortical cytoarchitecture [1]. This has motivated one of the central hypotheses of systems neuroscience: that cortex repeatedly instantiates a *canonical cortical microcircuit*. Under this view, common computational principles are reused across cortical areas, while specialization emerges through differences in connectivity, inputs, and learning rather than entirely different local operations [2]. Decades of work have proposed candidate canonical computations, including recurrent excitation and inhibition, normalization, gain modulation, recurrent amplification, and predictive coding [2, 3, 4, 5]. However, these computations have typically been evaluated against individual phenomena or local response properties. Identifying a common computational architecture that can account for the conserved organization of cortex across multiple levels, from individual neuronal responses to population structure and systems-level constraints, remains an open problem.

Computational modeling has made substantial progress in identifying principles of cortical computation. Task-optimized deep artificial neural networks (DANNs) [6] and their self-supervised counterparts [7] have emerged as powerful hypothesis spaces for understanding cortical responses across vision, audition, somatosensation, motor control, and language [8, 7, 9, 10, 11, 12]. In vision, convolutional neural networks (CNNs) are the best-studied of these models. Their local, hierarchical architecture, inspired by receptive fields in primary visual cortex, yields units that predict neural responses from early through late ventral visual cortex [6, 13, 14]. This success has made local, hierarchical computation the default account of how the ventral stream builds visual representations. But cortical circuits also show context-dependent changes in neuronal responses and contain long-range lateral connections [15, 16, 17], computations that standard CNNs do not include. How should these computations be captured in models, and what do they add to cortical organization?

Meanwhile, the *transformer* has become the dominant architecture in artificial intelligence (AI), supporting high task performance across language, vision, navigation, and reasoning [18, 19, 20, 21, 22]. Where CNNs build representations through local operations fixed by their architecture, transformers use learned multiplicative interactions to determine how information from different parts of an input influences one another. Information can therefore be routed flexibly across distant locations, with a much weaker inductive bias toward locality. These properties parallel two features of cortical circuits: context-dependent gain modulation and long-range lateral interactions. But because the transformer was not designed with cortical organization explicitly in mind, its apparent universal success raises a broader question: to what extent have evolution and machine learning discovered convergent or divergent computational solutions to the same underlying problem?

While early theoretical work linked transformer mechanisms to certain aspects of neural computation [23, 24], to thoroughly test the core empirical question at hand requires comparing the properties of many distinct candidate microcircuits—including, but not limited to CNNs and transformers. A truly correct account of a cortical microcircuit will need to match neural data better than a wide variety of alternatives, across multiple levels of brain organization. The computational tools and neural data needed to assess these comparisons at a breadth and depth demanded by a canonical microcircuit claim have only recently become available. Here we introduce a general strategy for making these comparisons. Our microcircuit-macrocircuit approach embeds competing microcircuits (e.g., CNNs, transformers, and others) within a shared hierarchical architecture with explicit cortical geometry, holding the macrocircuit fixed while varying the local circuit. This isolates the contribution of the microcircuit and lets us evaluate each candidate against a comprehensive, multi-scale account of what a canonical visual microcircuit must reproduce.

Applying this framework to the primate ventral visual stream yields a consistent answer across every level of analysis we tested: the transformer provides the most comprehensive account of ventral stream organization and function. At the single-unit level, transformers reproduce classical receptive fields, modulatory surround interactions, and the functional organization of lateral connectivity. At the population level, they develop hierarchical receptive fields with model depth, capture functional organization across early visual cortex and higher-order category-selective regions, and best predict held-out macaque neural responses. At the systems level, they combine strong task performance with reduced feedforward and lateral wiring costs. Mechanistic analyses indicate that this advantage arises primarily from learned multiplicative operations, rather than from access to long-range communication alone. These interactions route information between specific feature dimensions according to context, supporting contour detection and object-relation judgments. Together, these findings establish the transformer as a compelling candidate for a canonical cortical microcircuit and provide a general framework for inferring circuit principles across levels of biological organization.

### A microcircuit–macrocircuit strategy for evaluating canonical microcircuits

To systematically investigate which candidate microcircuit best accounts for core cortical signatures, we focus on the primate ventral visual system, whose organization and function have been characterized in great detail across multiple scales. We situate candidate microcircuits within a common Topographic Deep Artificial Neural Network (TDANN) “macrocircuit” (Fig. 1a), a framework previously shown to be effective in capturing functional organization in primate visual cortex and human language areas [25, 26]. TDANN generalizes standard hierarchical neural network models [27] by mapping each linear–nonlinear processing stage onto a two-dimensional cortical sheet and optimizing for both task performance and spatial smoothness, so that nearby units develop more similar tuning than distant ones.

**Figure 1.**
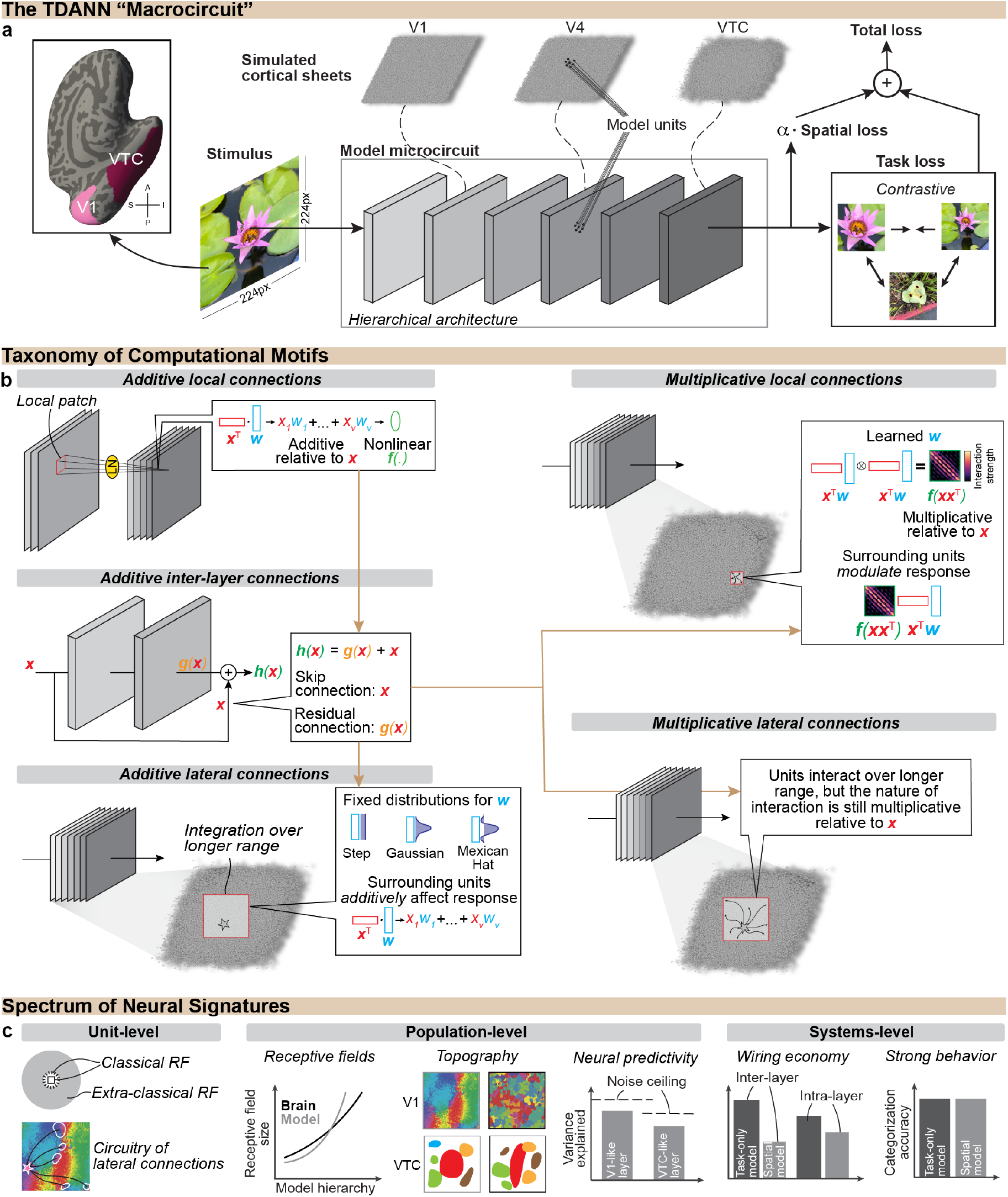
A macrocircuit–microcircuit strategy for evaluating candidate cortical computations. **a.** The Topographic Deep Artificial Neural Network (TDANN) provides the macrocircuit framework, specifying the system’s overall architecture and objective function. Models are hierarchical deep networks, with each layer represented as a simulated cortical sheet that assigns a spatial position to each unit. Models are trained on natural images to optimize task performance while satisfying a spatial constraint. **b**.0 Candidate microcircuits are instantiated within the macrocircuit. Each microcircuit candidate is defined by two properties of local computation: *how* information is combined, through additive or multiplicative operations, and *how far* information can travel, through local or longer-range connections. Arrows indicate motifs that are jointly implemented in leading AI architectures, illustrating how multiple motifs can be combined within a single microcircuit. Each candidate microcircuit is repeated across all model layers. **c**. Models are tested against a spectrum of neural signatures at three levels, spanning (i) unit-level, (ii) population-level, and (iii) systems-level comparisons.

With the macrocircuit fixed, we evaluate which local computation best instantiates the canonical microcircuit—the elementary operation performed by units within each layer, analogous to the local motifs governing how cortical neurons integrate their inputs. Each motif is defined along several dimensions: the spatial extent over which units communicate, and how they combine what they receive. Spatially, local connections keep wiring short but require many sequential steps to integrate information across distant spaces, while long-range lateral and skip connections shorten that path at the cost of additional wiring [15, 28]. Computationally, units may combine inputs additively, as in classical linear–nonlinear models, or multiplicatively, as in gain modulation and normalization [16, 4].

With these dimensions in mind, we instantiate and test a wide spectrum of motifs, including (Fig. 1b):

- *Additive local connections:* Units communicate only with nearby units and combine their inputs through *fixed additive operations*. Convolutional neural networks (CNNs), such as AlexNet [29], instantiate this motif.
- *Additive inter-layer connections:* Units communicate across layers through *additive longer-range shortcut connections*. Residual networks (ResNets) [30] extend CNNs with this motif.
- *Additive lateral connections:* Units communicate with other units within the same layer *additively* through *longer-range lateral connections*. Locally Laterally Connected Neural Networks (LLCNNs) [31] extend residual networks with this motif.
- *Multiplicative lateral connections:* Units communicate with other units within the same layer through *longer-range lateral connections*, with information integrated through *learned multiplicative operations*. Standard transformers [18, 19] instantiate this motif, alongside skip connections and local additive operations.
- *Multiplicative local connections:* Units communicate only with nearby units, but integrate information through *learned multiplicative operations*, as instantiated by Swin transformers [32].

#### A multi-scale battery of neural signatures

A canonical microcircuit’s brain alignment is judged by the consequences of its emergent signatures, which unfold across multiple scales of organization—from individual unit responses to behavior—and similarity at one scale may not guarantee similarity at another. We therefore evaluate each candidate with a series of detailed tests at each of three levels of neural scale (Fig. 1c):

1. At the *single-unit level*, we ask how units respond to visual information and what circuitry gives rise to these responses.
2. At the *population level*, we ask whether these local computations produce population-level organization: the hierarchy of processing across visual areas, maps of functional specialization, and predictive representations of held-out neural responses.
3. At the *systems level*, we ask whether the resulting system satisfies broader constraints faced by an organism: behavioral competence and efficient wiring. Together, these three levels provide an unprecedentedly stringent multi-scale test of each candidate microcircuit.

By evaluating outcomes at each level, we assemble a comprehensive picture of which microcircuit computation(s) correspond to core neural phenomenology.

### Local cortical computation and circuitry

#### Unit-level computation over the input

At the lowest level of analysis, a canonical microcircuit must reproduce the detailed response patterns of individual neural units or small groups of units. Much is known about how neurons in primary visual cortex (V1) respond to input. Responses within a localized region of visual space characterize V1 neurons’ classical receptive field. Responses are also modulated by contextual input from beyond this region, producing broader modulatory surrounds within their extra-classical receptive field (Fig. 2a; [34]). A correct model should capture both of these types of fine-grained response patterns. We therefore asked if the same detailed patterns are captured by any of the candidate microcircuits. First, we presented localized high-contrast sinusoidal gratings at different visual field positions and identified regions that evoked responses above baseline. Across most microcircuits, RF maps revealed localized excitatory and inhibitory subregions with small classical RFs, typically just above 1° in diameter (Fig. 2b). Second, we presented high-contrast gratings that expanded outward from the center of the visual field and measured two spatial response properties: the region over which responses grow with increasing stimulus size (the classical, or spatial summation, field), and the region beyond which responses asymptote following the peak (the extra-classical, or surround, field). Spatial summation fields were local across most architectures we tested and matched the classical receptive field sizes measured in macaque monkey V1, including transformers (Fig. 2c; one-sided Mann–Whitney *U* test, *U* =12, *n*=4 transformer and *n*=11 non-transformer models, *p*=0.143). Transformers were distinguished instead by the presence of surround fields, which closely matched those measured in macaque V1 (Fig. 2c; one-sided Mann–Whitney *U* test, *U* =0, *n*=4 transformer and *n*=11 non-transformer models, *p*=0.0028). Notably, transformers produced these broad modulatory effects despite using sparse rather than dense long-range connections (Fig. 2d).

**Figure 2.**
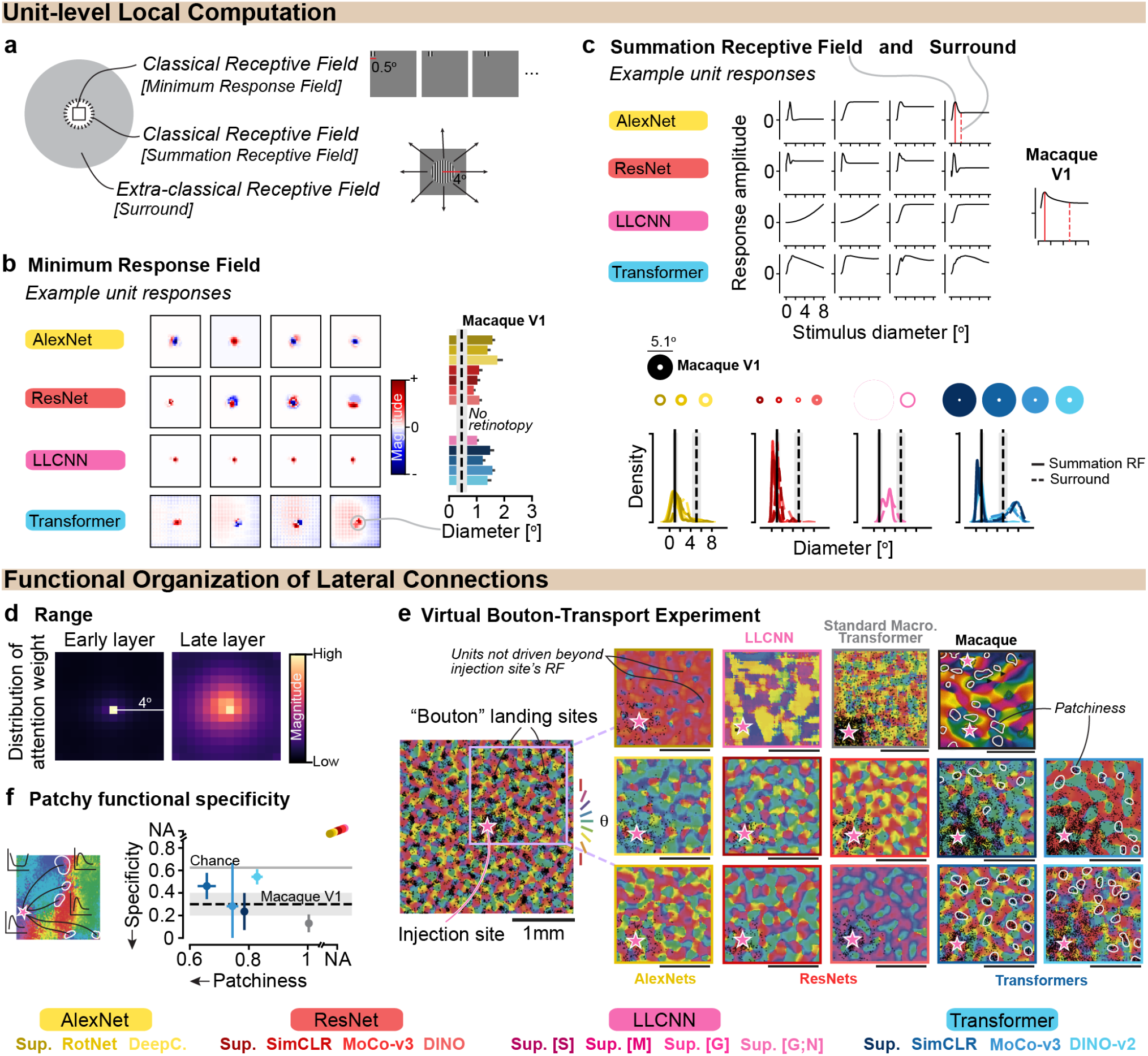
Evaluating details of local cortical computation and circuitry. **a**. Schematic of minimum response field (mRF), summation receptive field (sRF), and the surround for a V1-like unit. **b**. *Left*, example unit responses from sweeping a small high-contrast sinusoidal grating across the visual field. *Right*, minimum RF diameters for V1-like units across models. *Dotted black line*, macaque V1 estimate [33]. *Error bars*, 95% confidence interval (CI) over units. **c**. *Top*, example V1-like unit responses as a function of stimulus diameter in response to an expanding high-contrast sinusoidal grating. *Bottom*, distribution of sRF and surround diameters across models, with corresponding receptive field visualizations. *Black lines*, macaque V1 estimates [34]. *Error bars*, 95% CI over units. **d**. Distribution of attention allocation for early and late layers of an example transformer, computed over 1,024 randomly sampled ImageNet images and averaged over both images and attention heads. **e**. Virtual bouton landing sites (black symbols) shown over orientation preference maps for macaque V1 [15] and various models. Boundaries of clustered landing sites wherever identifiable were manually demarcated in white. **f**. Patchiness versus orientation specificity of landing sites. Points show the mean across 8 grating orientations. *Error bars*, standard deviation (SD) across the 8 orientations. *Dashed line and shaded band*, macaque V1 [15]. *Horizontal gray line*, chance (5/8, no relation between preferred and grating orientation). *Legend* —Models trained with different task objectives, grouped by microcircuit type.

#### Circuitry underlying long-range computation

At the level of circuitry, a canonical microcircuit should capture the organization of connections that produces these unit-level computations. In V1, contextual surround modulation is thought to arise from lateral connections that preferentially link neurons with similar feature preferences [34, 15]. Such connections were mapped experimentally by tracing the retrograde transport of biocytin from labeled boutons to their sites of origin. To test whether candidate microcircuits recapitulate the functional and spatial organization of these connections, we conducted a virtual bouton-transport experiment. We presented localized, high-contrast sinusoidal gratings at a fixed orientation to units in a V1-like layer, identified the most strongly modulated peripheral units outside the classical receptive field (the “landing sites”), and compared their preferred orientations with that of the stimulated unit. Transformers were the only microcircuit type that had stimulated units beyond the receptive field of the injection site (Fig. 2e). Moreover, these stimulated units exhibited a patchy arrangement, similar to the clustered horizontal connections observed in macaque V1 [15]. Quantifying this observation resulted in the TDANN-trained transformer models exhibiting patchy iso-orientation specificity close to the macaque V1 reference (Fig. 2f). Transformer models trained within the standard non-topographic macrocircuit, by contrast, showed landing sites that were diffusely and uniformly distributed across the cortical sheet, despite achieving comparable functional specificity. Thus, transformers uniquely recover both the functional *and* structural signatures of lateral connectivity when situated within the TDANN macrocircuit.

### Mid-level function and organization of cortex

#### Population-level receptive field hierarchy

We next examined how local computations combine across populations. A defining feature of the primate visual system is that populations at successive stages integrate information over increasingly large regions of visual space, creating a hierarchy from local to global representations [35]. Do candidate microcircuits, despite differing in their mechanisms of local computation, develop a comparable hierarchy of population receptive fields (pRFs). Several of the architectures we study—classic CNNs, ResNets, and LLCNNs—naturally display this property through stride and pooling operations, effectively building the pRF hierarchy into the architecture itself. Transformer microcircuits, however, can theoretically possess global pRFs at every layer due to their unconstrained attention mechanism (Fig. 3a).

**Figure 3.**
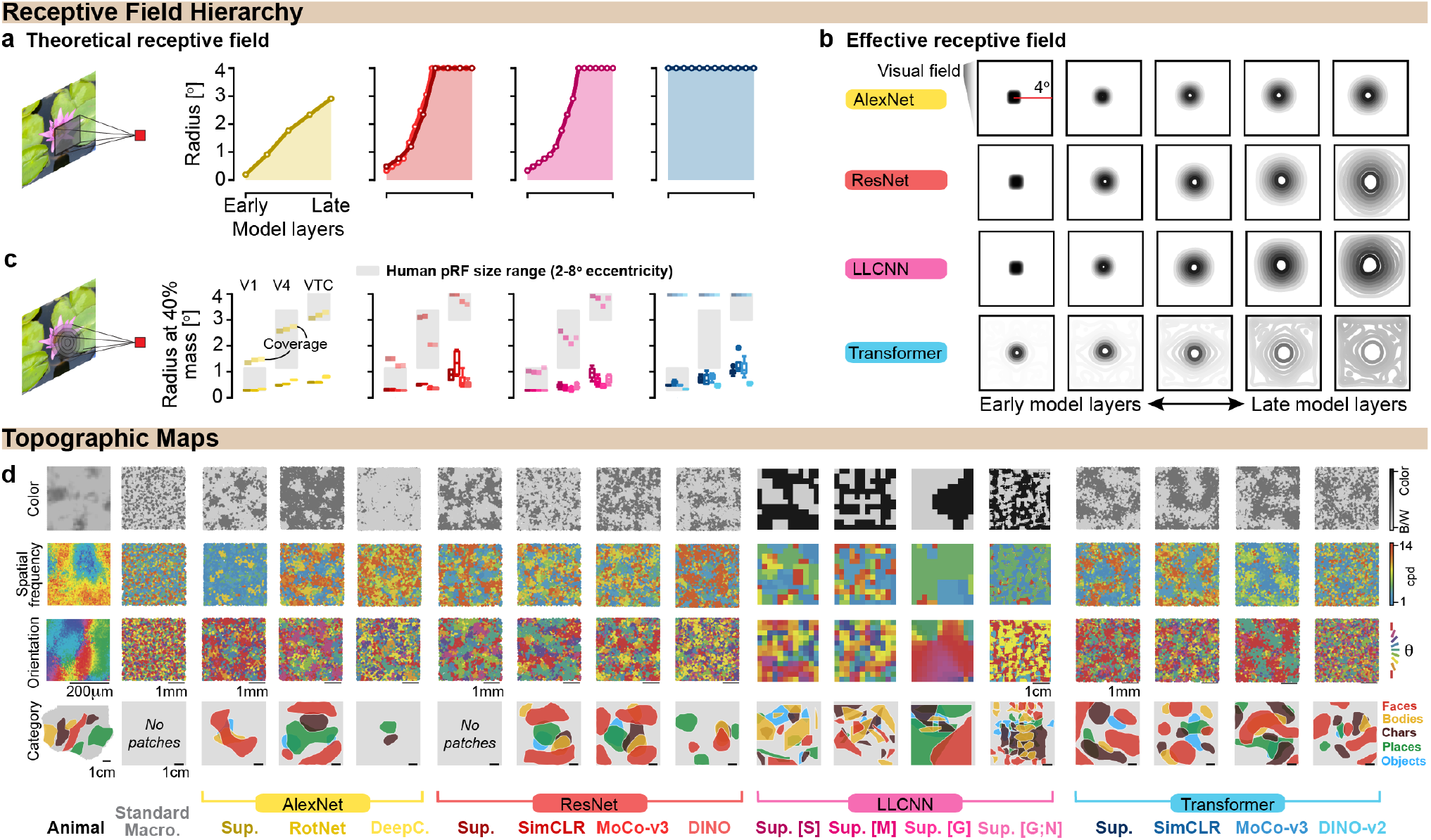
Evaluating population-level receptive fields and topographical organization. **a**.Theoretical maximum population receptive field (pRF) size across model layers. **b**. Kernel density visualizations of effective pRFs (EpRFs) for representative model layers. Maps are averaged over 1,024 randomly sampled natural images [36]. **c**. Radius enclosing 40% of EpRF mass for V1-like, V4-like, and VTC-like layers. Radius at 100% mass shown with lower opacity. *Gray*, human pRF size estimates [35]. *Error bars*, 95% CI over layers belonging to the corresponding brain area and 5 independent runs of the experiment. **d**. Color, spatial frequency, and orientation preference maps in macaque V1 [37] and most V1-like layer of various models. Smoothed category-selective maps in human VTC from NSD fLoc localizer [38] and in the most VTC-like layer of each model. See Suppl. Fig. S5 for quantification.

To determine whether this distinction persists after training, we measured effective population receptive fields (EpRFs) by computing the Jacobian of unit responses with respect to the input image to characterize the portion of the visual field that modulates those responses, including both the classical RF and surrounding contextual influences [39]. We find that trained transformers develop a local-to-global EpRF hierarchy despite their theoretically global architecture (Fig. 3b; also see Suppl. Fig. S1). We quantify this by measuring the *radius enclosing 40% of EpRF mass*—analogous to the standard deviation of a Gaussian pRF profile. Radius increases monotonically across V1-like, V4-like, and VTC-like layers in all architectures (Fig. 3c; one-sided one-sample t-tests of across-region slopes against zero, all *p*<1.3 × 10^−6^), suggesting that all microcircuit taxa develop receptive-field organization broadly consistent with the ventral visual hierarchy.

#### Multi-areal cortical topography

A canonical microcircuit should reproduce not only how units compute over the input, but also how those learned features organize into cortex’s mid-level architecture. In V1, neurons are selective for features such as orientation, spatial frequency, and color, and these preferences form structured topographic maps across the cortical sheet [40, 41, 42]. Across the most V1-like layers of our models, nearly all microcircuits embedded within the TDANN macrocircuit develop orientation, spatial frequency, and color maps resembling those observed in macaque V1 (Fig. 3d), with LLCNNs using weight sharing the primary exception, showing substantially coarser spatial organization. Further along the visual hierarchy, category-selective responses in ventral temporal cortex likewise form spatially organized patches for faces, bodies, words, and places [43], and TDANN-trained microcircuits reproduce these patches as well (Fig. 3d). Critically, these organizational signatures are not explained by task training alone (“standard macrocircuit” models in Fig. 3), but require the spatial constraints imposed by the TDANN macrocircuit. Quantitative analyses confirm the correspondence between model-derived topographic maps and those observed in cortex (Suppl. Fig. S5).

#### Neural response pattern similarity

Beyond reproducing cortical organization and its underlying circuitry, a successful model should predict the fine-grained pattern of neural responses evoked in each neuron by individual stimuli. We therefore evaluate model unit response pattern similarity to real neural responses from the THINGS Ventral Spiking Dataset (TVSD) [44], a large-scale multi-area electrophysiology dataset recorded at millisecond resolution while two macaques viewed over 25,000 natural images. We compare model and neural responses in both directions using the Inter-Animal Transform Class (IATC) approach [45], which tests whether a model’s representations can substitute for another animal’s in the population and thereby offers both high predictivity and high systems-identification specificity. Mappings between the two systems are estimated using bidirectional linear ridge regression [45, 46], in which models are required both to predict real neural responses under regression reweighting (the standard “model-to-brain” direction [6]), as well as be predicted *by neurons* (the reverse “brain-to-model” direction). By requiring per-neuron stimulus-by-stimulus predictivity in both directions, this methodology imposes a strong test on model-brain neural response pattern similarity [45].

The model-to-brain mapping results showed that all models explained substantial neural variance, with limited separation between microcircuit classes (Fig. 4a). This non-separating result is consistent with earlier results [25]. However, we also found that reverse mappings from brain-to-model provided a more stringent test: microcircuits embedded within the TDANN macrocircuit consistently achieved higher neural predictivity than their non-topographic counterparts across brain areas (Fig. 4b). The reverse regression test was able to differentiate between microcircuits, with the transformer circuit exhibiting both the strongest brain-to-model alignment, and the strongest responsiveness to topographical constraints. Layer-wise neural predictivity for all brain areas is reported in Suppl. Fig. S6. Consistent with this result, topographic models exhibited lower intrinsic dimensionality [47] than non-topographic models (Fig. 4c; layer-wise results in Suppl. Fig. S7), showing that the topographic constraint compresses population representations toward the empirical dimensionality of the true neural representation. Topographic transformers maintained high neural alignment despite this compression, implying that this microcircuit motif is particularly effective at supporting compact yet expressive representations.

**Figure 4.**
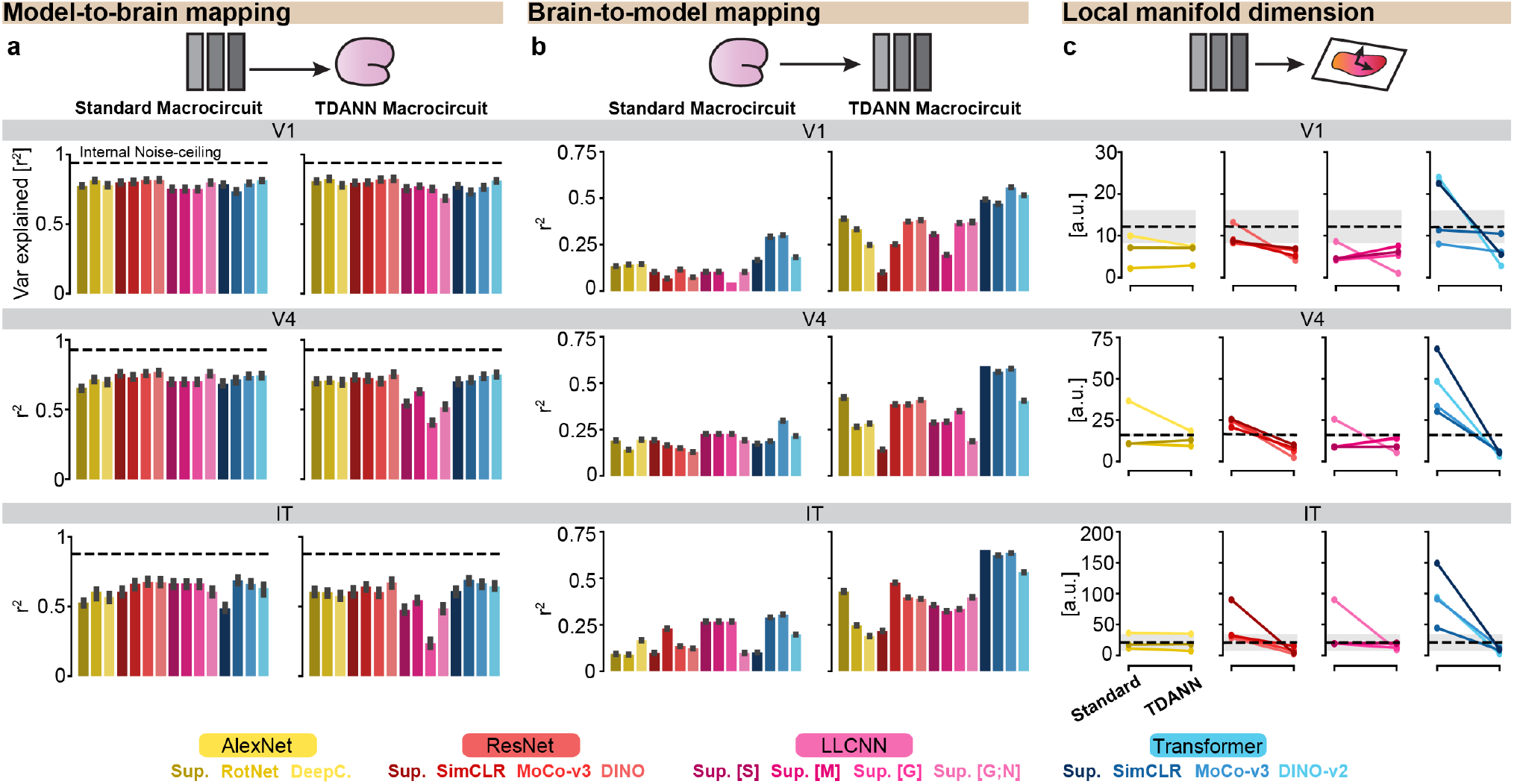
Evaluating brain predictivity and intrinsic dimensionality of functional responses. **a**.Median variance explained by a linear mapping from the most V1-like, V4-like, and IT-like model layers to multi-unit spiking activity recorded in V1, V4, and IT from two monkeys [44]. *Black line*, Spearman–Brown-corrected split-half consistency. *Error bars*, 95% CI over electrodes. **b**. Same as **(a)**, but for the reverse mapping from brain activity to model responses. The noise ceiling is 1.0, because model responses are deterministic across trials. *Error bars*, 95% CI over units. **c**. Intrinsic dimensionality of the most V1-like, V4-like, and IT-like model layers, computed over 1,024 images randomly sampled from the training images shown to the two monkeys. *Black line*, mean dimensionality of monkey neural responses, with shading indicating variability across monkeys. *Standard Macrocircuit*, models trained using only the task objective; *TDANN Macrocircuit*, models trained with the TDANN objective. For each model, V1-like and VTC/IT-like layers were defined as the layers whose responses best predicted neural activity in the corresponding brain region (Suppl. Fig. S6).

### Systems-level cortical constraints

#### Feedforward and lateral wiring costs

A strong constraint on the emergence of cortical topography is wiring economy: because neural tissue is spatially embedded, shorter connections reduce material, metabolic, and developmental costs [48, 49]. We thus test whether candidate microcircuits, when embedded within the TDANN macrocircuit, reduce wiring costs both between and within processing stages. Feedforward wiring captures projections between successive processing stages (Fig. 5a left), whereas lateral wiring captures interactions within a cortical area (Fig. 5b left). Because LLCNN connectivity is fixed a priori, changes in its lateral wiring reflect manually prescribed connection lengths rather than learned spatial organization. Transformers, in contrast, learn their within-layer interactions through attention, providing an opportunity to test whether topographic optimization jointly reduces feedforward and lateral wiring within a common modeling framework.

**Figure 5.**
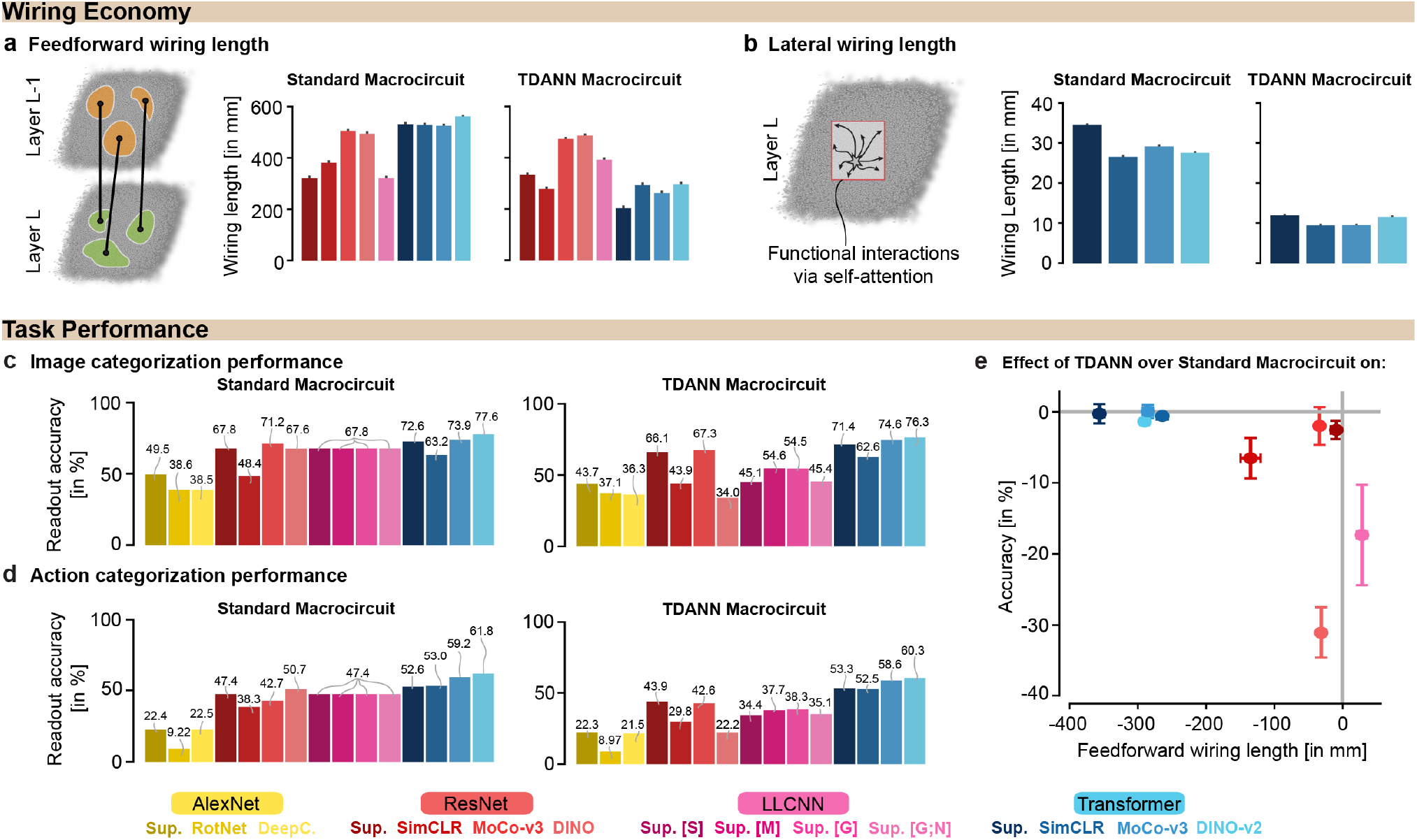
Evaluating systems-level constraints of wiring economy and task performance. **a**. Feedforward wiring length for each model’s most VTC-like layers, computed functionally over 1,024 randomly sampled ImageNet images. *Error bars*, 95% CI over cortical neighborhoods and input images. AlexNet and LLCNN models with weight sharing are omitted due to insufficient depth or coarse retinotopy. **b**. Lateral wiring length for each model’s most VTC-like layers. *Error bars*, 95% CI over model units. LLCNNs are omitted because their lateral wiring length is fixed during training. **c**. Top-1 image categorization accuracy on the ImageNet validation set under a linear readout. **d**. Top-1 action categorization accuracy on the Kinetics400 [50] validation set under a linear readout. **e**. Relationship between the reduction in feedforward wiring length and the reduction in task performance under different macrocircuit types. *Error bars*, 95% CI over the two tasks along the y-axis, and 95% CI over neighborhoods and images along the x-axis.

We first measured feedforward wiring as the total length of connections between adjacent model layers. Most topographic models exhibited shorter feedforward wiring than their non-topographic counterparts, with the exception of ResNet Supervised and LLCNN (Fig. 5a right; one-sided Wilcoxon signed-rank tests, *n*=200 paired shift directions and images sampled, *p*<0.001). We next measured lateral wiring, treating attention weights as a proxy for the strength of within-layer interactions and computing the distance-weighted cost of the resulting interaction graph.

Spatial optimization shifted transformer interactions toward shorter distances, resulting in substantially shorter lateral wiring in topographic transformers than their non-topographic counterparts (Fig. 5b right; one-sided Wilcoxon signed-rank tests, *n*=3072 paired units and attention heads, *p*<0.001). The layer-wise breakdown of wiring costs is provided in Suppl. Fig. S8.

Together, these results show that topographic transformers simultaneously reduce wiring costs between and within processing stages. Thus, the same microcircuit that reproduces low- and mid-level cortical functional organization also satisfies a fundamental high-level physical constraint thought to shape the architecture of the visual system.

#### Task performance

The visual system must balance the physical costs of wiring with the demands of sophisticated visual behavior [51, 52]. Spatial constraints can improve biological organization in neural networks, but can come at a cost to task performance [25, 31]. An optimally effective microcircuit would maintain behavioral competence while satisfying this pressure for wiring economy. To assay visual behavioral competence, we evaluated performance on two challenging naturalistic visual tasks that have been extensively used to benchmark primate visual system models in recent works [53, 54]: 1000-way object recognition on ImageNet [36] and 400-way action recognition on Kinetics-400 [50]. Under the standard non-topographic macrocircuit, transformers performed comparably to or somewhat better than other microcircuit classes across both tasks (Fig. 5c–d, left panels; one-sided Mann–Whitney *U* test, *n*=4 transformer and *n*=11 non-transformer models; 2 tasks; *U* =37, 44; *p*=0.1078, 0.0085, respectively).

Critically, transformers were substantially less susceptible than alternative microcircuits to performance loss under the topographic TDANN macrocircuit (Fig. 5c–d, right panels). Moving from the standard to the TDANN macrocircuit did not substantially degrade transformer top-1 accuracy (median Δ=-0.6 percentage points, worst case −1.4 pp).

Non-transformer models dropped more (median −7.2 pp), with the largest declines in LLCNNs and ResNet50 DINO (up to −33.6 pp). This is consistent with prior work showing a tradeoff between wiring-length optimization and task performance [25]. Transformers therefore maintain high behavioral performance while reducing both feedforward and lateral wiring costs—the only microcircuit class to satisfy both high-level organismic demands simultaneously (Fig. 5e).

### Robust local multiplicative circuits for relationship detection

The apparent convergence of transformers with cortex across all levels of analysis raises two related questions: What mechanistic component(s) of the transformer microcircuit drives convergence? And what is the specific functional role of that mechanism?

#### Avoiding dense long-range communication

The microcircuit taxonomy naturally separates motifs along two key dimensions—the spatial extent over which units communicate, and the nature of the computation they perform. Transformers differ from CNNs, ResNets, and LLCNNs along both dimensions, combining learned multiplicative interactions with substantially longer-range communication (Fig. 6a). The first key mechanistic question is thus: do measures of cortical convergence benefit from transformer’s extended communication range, the multiplicative computation type, or a combination of both factors?

**Figure 6.**
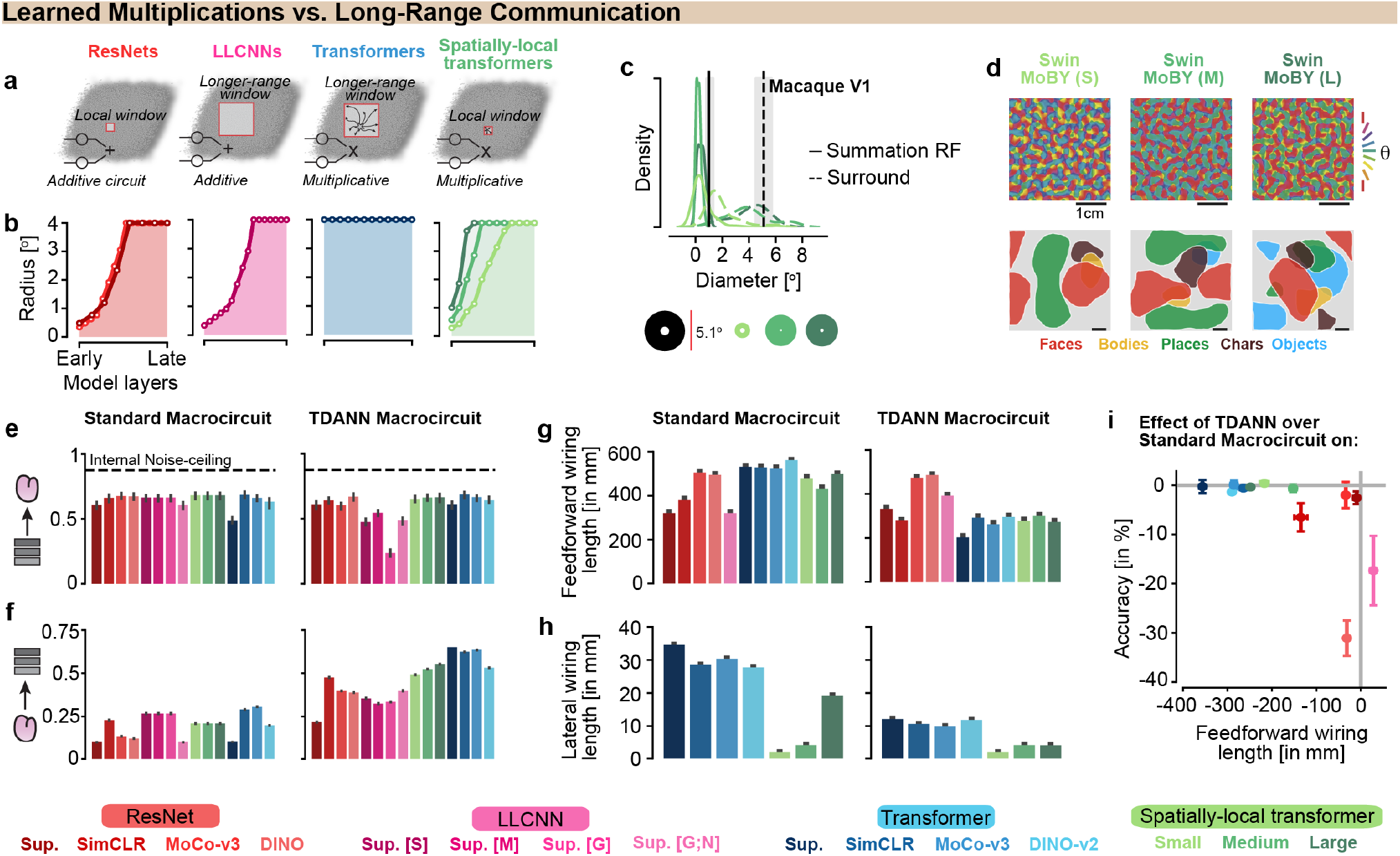
Multiplicative operations, rather than dense connectivity *per se*, support transformer–brain convergence. **a**. Schematic of the microcircuit motifs for ResNets, LLCNNs, standard transformers, and the local transformer variants. **b**. Theoretical pRF size across model layers. **c**. Summation RF and surround size for model V1 units. **d**. Orientation preference maps and category selective maps for different models. **e**. Model-to-brain predictivity, measured using explained variance, between most IT-like model layers to macaque IT responses. **f**. Brain-to-model predictivity, measured using explained variance, from macaque IT responses to most IT-like model layers. **g**. Feedforward wiring length in the most VTC-like model layers. **h**. Lateral wiring length in the most VTC-like model layers. **i**. Relationship between the reduction in feedforward wiring length and the reduction in task performance under different macrocircuit types. Plotting conventions, including definitions of error bars, follow those of the previous figures.

To address this question, we instantiated an additional class of microcircuits that retain the computational features of multiplicative attention while restricting the spatial extent of interactions to match that of inherently local models like CNNs. Such spatially-restricted transformer-like architectures were introduced with the Swin transformer architecture [32].

We evaluated three spatially-local transformer variants with small, medium, and large attention windows against the full battery of cortical signatures. Varying window size provides a controlled continuum from CNN-like locality to unconstrained global attention without changing the underlying computation (Fig. 6b), allowing communication range to be manipulated independently of computation type. All three performed comparably to transformers with unrestricted global attention across these assays. They developed small summation receptive fields flanked by modulatory surrounds, whose size varied with the attention window (Fig. 6c). At the population level, they reproduced functional maps in V1 and VTC (Fig. 6d) and achieved strong bidirectional neural predictivity (Fig. 6e–f). Strikingly, spatially local transformers minimized both feedforward and lateral wiring costs without compromising task performance (Fig. 6g–i). These findings suggest that learned multiplicative computation, rather than extended spatial communication *per se*, is the primary driver of transformer–cortex convergence.

#### Privileged multiplicative routing along privileged axes supports relational readouts

What specific computational advantage(s) does the local multiplicative circuit endow the network? Wiring economy creates a fundamental tension: reducing physical connectivity lowers wiring cost, but purely local additive circuits must build long chains of computation to integrate information across distant elements of a scene. Multiplicative interactions provide a simple way around this constraint. A clue comes from the study of large language models, in which linear readouts can be shown to readily recover properties of individual entities, whereas relationships between entities emerge through sparse token-to-token multiplicative interactions [57, 58]. We tested whether the same principle holds in vision: do learned multiplicative interactions provide a general mechanism for representing relationships between distant elements without requiring dense long-range connectivity?

To evaluate this idea, we adopted the recently-introduced concept of *privileged axes* [59]. Privileged axes are special directions in representational space that consistently arise at the single-unit level, across independently-trained networks [59, 60]. We reasoned that, if they arose in transformer networks, privileged axes would be the natural “operands” for relational interactions. We empirically tested the existence of such axes by matching units in the GeLU-MLP layers of multiple independently trained transformer models. Privileged axes were reliably identified throughout the model hierarchy (Fig. 7a left; one-sided Mann–Whitney *U* test against chance, *p*=10^−5^). To understand the contents of the privileged axis, we probed their response properties on datasets of stimuli constructed to capture contour integration [55] and object relationships [56]. This revealed a progression in the content of the privileged axes: orientation emerged as a privileged axis in early layers, consistent with previous results [59], while object shape and color emerged in later layers (Fig. 7a right). Thus, the candidate “operands” for multiplicative attention computation are not arbitrary features, but structured and reproducible elementary properties of the visual representation.

**Figure 7.**
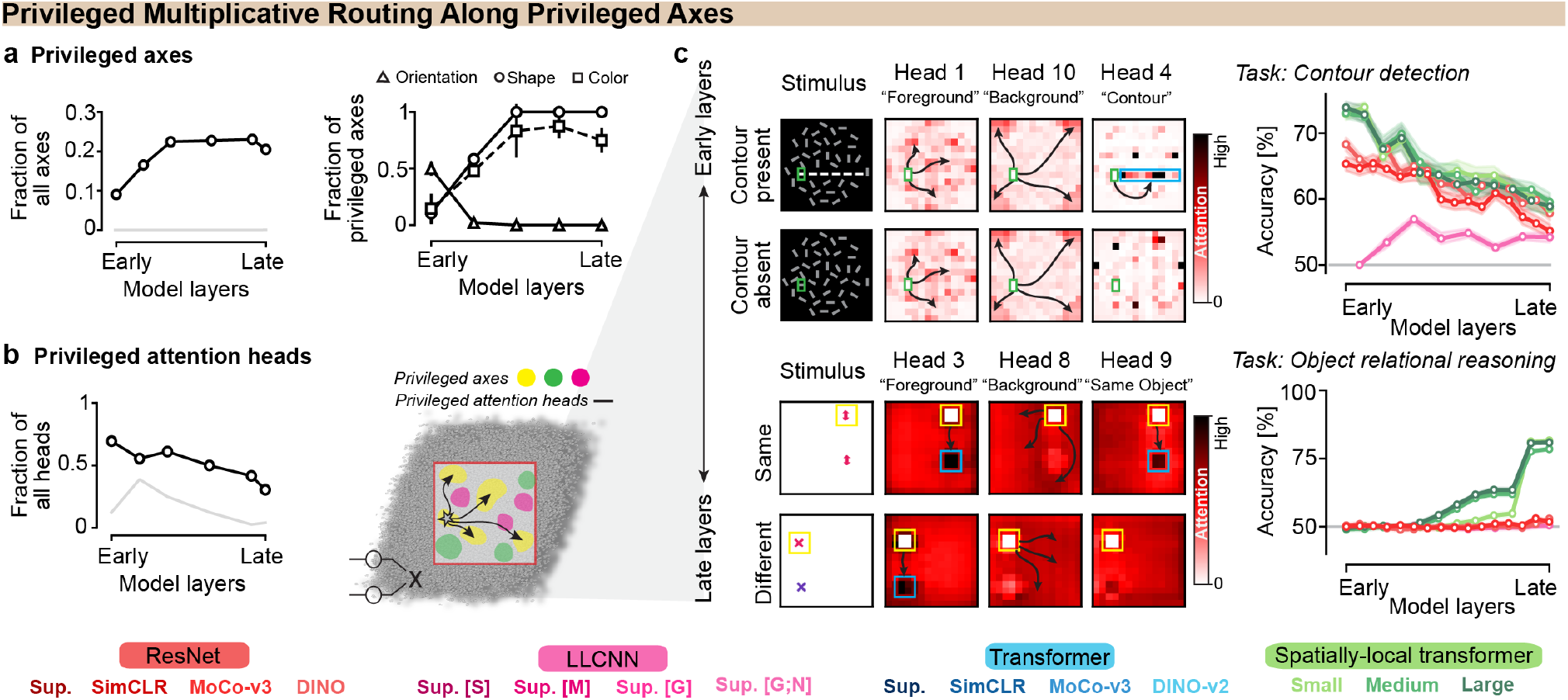
Relational readouts are subserved by privileged multiplicative routing along privileged representational axes. *a*. *Left*, fraction of all transformer model axes that are privileged. *Chance*, repeating the method for a permuted image order. *Right*, fraction of privileged axes explaining at least 10% of the variance in any property. *Error bars*, SD across three model seeds. **b**. *Left*, fraction of all attention heads that are privileged. *Chance*, repeating the method for a permuted image order. *Error bars*, standard deviation (SD) across three model seeds. *Right*, schematic of privileged attention heads multiplicatively routing information over privileged axes in the transformer. **c**. *Top left*, example stimuli from a contour-detection task [55] and corresponding attention heads routing information from tokens within one contour segment (green) to the rest of the image. *Bottom left*, example stimuli from an object relational reasoning task [56] and corresponding attention heads routing information from tokens of one object (yellow) to the rest of the image. *Right*, linear readout accuracy for contour detection and object relational reasoning across model layers. Local transformers and ResNets are parameter-matched to ensure fairness of performance comparison.

We next asked what types of multiplicative interactions the transformer attention implemented relative to the privileged axes. Multiplicative attention in transformers is mediated by “heads”—units that express attentional “routing patterns” linking pairs of input MLP axes [61]. Recent mathematical theory of neural networks [60] predicts the existence of *privileged attention heads* complementary to the privileged axes—that is, specific routing patterns that arise robustly across multiple independently-optimized model instances. We first empirically confirmed the existence of such privileged attention heads throughout the network hierarchy (Fig. 7b left; one-sided Mann–Whitney *U* test against chance, *p*=10^−5^).

We then analyzed in greater detail what these robustly-appearing multiplicative routing patterns actually do. We found that, across the model hierarchy, privileged attention heads selectively linked tokens according to similarity on the privileged visual axis dimensions (Fig. 7b right). These multiplicative routing patterns are highly interpretable. Routing patterns in early network layers preferentially link orientation-selective axes (Fig. 7c, top left), consistent with results on non-classical receptive fields described earlier. As a consequence, reliably-appearing heads effectively compute a contour-integration signal [55]. In later network layers, routing patterns also compute same-different signal across the object properties captured by the preferred axes—in this case, object identity (Fig. 7c, bottom left). Visualizations of all attention heads and their routing patterns across both tasks are provided in Suppl. Fig. S9.

Critically, these interactions were functionally consequential. Linear readouts from early transformer layers support strong contour detection readouts (Fig. 7c, top right). Similarly, later layers supported relational reasoning between objects (Fig. 7c, bottom right). In both cases, architectures without learnable multiplicative components performed poorly or failed altogether on these relationship extraction tasks. Together, these results reveal how multiplication enables efficient relational computation: privileged heads route information along privileged axes, and their multiplicative interactions make relationships explicitly available for readout, without requiring dense long-range connectivity.

## Discussion

A central question in systems neuroscience is whether the diverse functions of cortex can be supported by a common set of local computations, often framed as a *canonical cortical microcircuit*. Here we show that the transformer, the dominant architecture in machine learning, has many of the hallmarks one would expect of such a computational primitive. We embedded a taxonomy of candidate microcircuits within a shared topographic macrocircuit, so that each could be evaluated across multiple levels of cortical organization while the surrounding architecture was held fixed. In the primate ventral visual stream, the transformer most consistently met the criteria expected of a canonical cortical microcircuit, capturing single-unit computation, population-level functional organization and response properties, and systems-level wiring and task demands. Mechanistically, this advantage traces primarily to learned multiplicative interactions, in which privileged attention heads route relational information between specific privileged representational axes without requiring dense connectivity. Together, these findings suggest that the transformer and the mammalian cortical microcircuit may reflect similar solutions to the problem of building intelligent systems under biological constraints.

The advantage of multiplicative interactions lies not in what a network with multiplicative routing primitives can represent, but in how efficiently it represents relationships. The universal approximation theorem guarantees that sufficiently large linear–nonlinear (LN) networks can approximate any continuous function in principle [62]. Yet much of representational neuroscience has focused on which architectures best predict cortical responses, without distinguishing predictive accuracy from the computations underlying it [54, 63]. Given sufficient capacity, models trained on the same data and objective may learn similarly predictive input–output mappings despite differences in their computational operations. This offers a potential explanation for why large-scale comparisons have found similar neural predictivity across architectural families [63]. Our results highlight an important distinction that predictive accuracy alone can obscure: representational expressivity does not imply computational efficiency. Additive LN networks may require substantially greater depth, width, or wiring to represent relationships that circuits incorporating multiplicative operations can capture more efficiently. This distinction is particularly consequential for biological systems, where circuit structure constrains both learning and implementation. Multiplicative operations may therefore confer an advantage not in what can be represented, but in how efficiently relational computations can be learned and implemented.

Our results do not specify a specific low-level mechanistic implementation of multiplicative operations in brain-cellular “hardware”—though our spatially-registered topographical evaluations provide strong convergent evidence that this implementation, if it is indeed present, should exist at the scale of a single neuron or small group of neurons. Multiplicative-like relational computation can be realized through different biological substrates, a distinction others have drawn independently in mapping transformer operations onto laminar cortical organization [24]. Candidate mechanisms extending beyond local linear filtering include the two-layer neural network interpretation of pyramidal cells [64] and neuromodulatory gating by glial cells [23]. Rigorous empirical evaluation of these proposals remains an important direction for future work.

From an evolutionary biology standpoint, constraints on ecologically relevant behavior and biophysical connection length are fundamental. Theories of ‘wiring optimization’, ‘save wire’, and ‘laws of protoplasmic economy’ [65, 66, 48] all arise from the need to use metabolic and material resources efficiently in service of functional anatomy, and been proposed as a general organizing principle across scales [67, 68]. The TDANN macrocircuit operationalizes these two constraints directly, and our results reinforce its value as a framework for understanding cortical organization across scales. Importantly, this framework enables a new mode of inference: by identifying which microcircuit types satisfy both constraints, we can reason backward from systems-level organization to underlying circuit structure. The finding that multiplicative operations provide an effective solution to these constraints suggests that similar computational motifs may have been favored during the evolution of cortical circuits.

A key limitation of our work is that several micro- and macrocircuit classes fall outside our direct comparison. At the microcircuit level, All-TNNs [69] eliminate convolutional weight sharing, allowing spatially distinct weights while enforcing local smoothness. Although they reproduce topographic organization and center–periphery structure in V1 and VTC, this organization is much coarser and makes some benchmarks, such as pinwheel density, difficult to interpret, precluding direct comparison. We therefore include a not weight-shared LLCNN variant as a partial proxy. At the macrocircuit level, TopoNet [70] suppresses high-frequency variation in filter weights and has reproduced mid-level VTC organization with both CNN and transformer microcircuits, while the interactive topographic network (ITN) [71] explicitly optimizes wiring cost alongside task performance. Both frameworks have primarily been evaluated in higher visual cortex, however, leaving open whether they can reproduce the fine-grained tuning and spatial organization observed in V1.

It is worth considering the broader pervasiveness of the transformer architecture. Recent work has shown that a language transformer trained under the TDANN framework with a next-word prediction objective captures key aspects of the functional organization of the human language system [26]. It is striking that the same underlying circuit motif can account for functional organization constraints in two seemingly distinct domains: vision and language. This convergence suggests that a task-agnostic self-supervised spatial constraint combined with an additive plus multiplicative microcircuit may reflect a more general computational principle of cortex. Such a framework could potentially extend beyond language and vision, providing a unified architecture for modeling and predicting the functional organization of sensory, motor, and cognitive systems.

## Methods

Some evaluation metrics were adopted from [25]. Here, we describe only the metrics that were newly introduced or independently implemented in the present work.

### Model specification

#### Model units

What constitutes a model unit? In convolutional architectures such as AlexNet, the rectified output of each convolutional layer provides a natural definition of a model unit. In ResNets, following [25], model units are defined as the outputs of residual blocks, which likewise terminate in a rectification operation. In LLCNNs, each convolutional layer produces a population of model units. Transformer-based architectures are less straightforward. At every layer, the model represents the image as a set of learned token embeddings, and either the outputs of the transformer layer itself or intermediate representations within the layer could plausibly serve as model units. This raises a key question for topographic modeling: which representation should be associated with cortical responses? To address this question, we focus on the attention module, the component through which token–token interactions occur. We consider two candidate definitions. The first defines model units as those whose representations enter the attention module of the next layer, producing responses before lateral interactions have been incorporated. The second defines model units as those whose representations form the output of the attention module, producing responses after integrating information from other tokens within the layer. Empirically, the latter produces substantially more accurate topographic models. Accordingly, all subsequent analyses adopt attention outputs as the definition of model unit responses (see Suppl. Fig. S3 for results using the alternative definition).

#### Model size and task objectives

We evaluate a diverse set of model architectures and training objectives. AlexNet [29] was trained using three task objectives: supervised learning, RotNet [72], and DeepCluster [73]. ResNet-18 [30] was trained with supervised learning and SimCLR [74], while ResNet-50 was trained with MoCo-v3 [75] and DINO [76]. LLCNN models [31] were trained in a supervised manner with three different lateral connectivity profiles—Gaussian, Mexican-hat, and step-function weighting—as well as a variant combining Gaussian-weighted lateral connections with a locally connected layer that does not share weights across spatial locations. Importantly, LLCNN models were trained on the task loss alone as proposed by their authors [31]. Finally, Transformer-Base models [19] were trained using supervised learning, SimCLR, MoCo-v3, and DINO-v2 [77]. All models were trained on a natural image diet (ImageNet [36]). Abbreviations used throughout the paper—*Sup*: Supervised, *DeepC*.: DeepCluster, *LLCNN [S]* : Step-function weighted lateral connections, *LLCNN [M]* : Mexican-hat weighted, *LLCNN [G]* : Gaussian weighted, *LLCNN [G;N]* : Gaussian weighted without weight sharing.

#### Position assignment

Positions for CNN units were assigned following the procedures of [25] and [31]. For transformers, each token embedding was initially assigned a retinotopic position based on the image patch from which it was derived, and tokens were scattered across a two-dimensional simulated cortical sheet accordingly. Prior to training, we applied a swapping optimization (“swapopt”) procedure similar to that of [25], which reorganizes unit positions in response to simple sinusoidal gratings to establish an initial topographic layout. Neighborhood widths used for the spatial loss were taken directly from [25].

#### Training procedure

All models were trained using the training configurations provided by the authors of the respective task objectives. Spatially optimized models were trained in the same manner, with the addition of a spatial loss term to the task objective. Unless otherwise noted, the spatial loss was weighted by *α* = 0.25 across all model variants, corresponding to the optimal value reported by [25]. We additionally verified the robustness of this choice by sweeping over a range of *α* values for a representative transformer model and observed qualitatively similar results (Suppl. Fig. S4).

### Receptive field analysis

#### Theoretical population receptive field size

For layer *l* and an assumed input of size 224×224, the theoretical pRF size was calculated as:

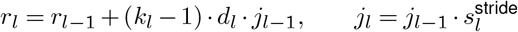

initialized at *r*_0_ = 1, *j*_0_ = 1, *s*_0_ = 0.5, where *k*_*l*_, *d*_*l*_, *p*_*l*_, and 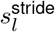 are the kernel size, dilation, padding, and stride of layer *l*.

#### Effective population receptive field globality index

For a given layer *i*, we first obtained the gradient of a scalar summary of that layer’s central activations with respect to the input pixel space. Specifically, let 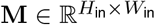 denote the resulting gradient map, computed as the absolute value of 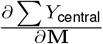, where *Y*_central_ is the sum of absolute activations at the spatially central positions of the feature map—defined as a *s*×*s* region centered on the feature map, where *s* is a layer-specific scale factor chosen to subtend the same relative spatial extent across architectures. To summarize this map as a scalar measure of locality, globality index was defined as:

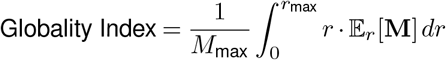

#### Minimum response field size

mRFs were mapped by presenting small circular sine grating patches (0.5° diameter, 14px radius) at each position on a regular grid (spacing of 7px) covering the full 224 224 image. Each patch was rendered at 4 orientations (0°, 45°, 90°, 135°), 4 spatial frequencies (18.6, 31.6, 57.8, 96.9 cpd), and 4 phases (0, *π/*2, *π*, 3*π/*2), yielding responses that were averaged across all orientation, spatial frequency, and phase conditions at each grid position to produce a spatial response map per unit. Analysis was restricted to units at spatially central positions of each feature map. Baseline-subtracted responses were normalized to match the median peak firing rate of V1 neurons reported by [78]. The spatial response map was thresholded at 5% of its peak response, and only the single contiguous suprathreshold region containing the driven central grid position was retained; units with no suprathreshold response at this position were excluded. mRF diameter was estimated by counting the number of grid positions within this contiguous region and back-computing the diameter of a circle with equivalent area:

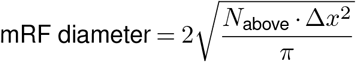

where *N*_above_ is the number of suprathreshold positions within the contiguous region and Δ*x* is the grid spacing in degrees.

#### Summation receptive field and surround size

Expanding circular apertures were applied to full-field sine gratings (8 orientations, 8 spatial frequencies, 4 phases, 2 color conditions) at 28 aperture diameters ranging from 0° (blank gray baseline) to 8°. Responses were baseline-subtracted and restricted to orientation-selective units at central feature map positions, where orientation selectivity was defined as circular variance < 0.6 and a positive response at the preferred orientation at the largest aperture. Size tuning curves were constructed by averaging responses across stimuli at each unit’s preferred orientation as a function of aperture diameter.

Summation RF size was defined as the aperture diameter at peak response, identified as the first local maximum in the size tuning curve. The surround size was defined as the smallest diameter beyond the peak at which the response remained within 10% of the asymptotic value for all larger diameters.

### V1 and VTC topography benchmarks

#### Pinwheel density

Orientation preference maps were smoothed using a Gaussian kernel (width = 1.5 mm, stride = 0.15 mm) and pinwheels were detected via winding numbers computed over the resulting map. For each interior pixel, the sum of circular increments around its 3 × 3 neighborhood was computed; a winding number of 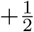 or 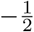 indicates a positive or negative pinwheel, respectively. Candidate pinwheel pixels were thresholded at |*w*| > 0.3 and additionally required to have local circular standard deviation < 0.3, to exclude spurious detections in noisy regions. Contiguous clusters of fewer than 2 pixels were discarded, and the centroid of each surviving cluster was taken as the pinwheel center.

Pinwheel density was normalized by hypercolumn spacing Λ, estimated from the orientation map’s power spectrum. The smoothed complex orientation map *e*^2*iθ*^ was Fourier transformed, and a Gaussian was fit to the radially averaged power spectrum to identify the peak spatial frequency *f* ^∗^; column spacing was then Λ = 1*/f* ^∗^, converted to mm using the known spatial extent of the tissue. Pinwheel density was reported as *ρ* = *N/*(*A/*Λ^2^), where *N* is the total pinwheel count and *A/*Λ^2^ is the tissue area in units of hypercolumn spacing squared.

#### Circular variance

CV was computed following [78]. Orientation tuning curves were obtained by averaging responses across spatial frequency, phase, and color conditions, then computing:

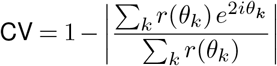

over 8 equally spaced orientations *θ*_*k*_ ∈ [0°, 157.5°]. Tuning curves were shifted to a non-negative baseline prior to CV computation.

### Functional organization of lateral connections

To assess the orientation specificity of model lateral connections, we presented small circular gratings (0.5° aperture) at the image center at 8 orientations (*θ* ∈ {0°, 22.5°,…, 157.5°}). We averaged responses across spatial frequency, phase, and color, and subtracted the mean response to a blank gray screen. This gave each unit’s modulation *M*_*i*_(*θ*). Each unit’s preferred orientation was the orientation eliciting the largest mean response to full-field (8°) gratings.

#### Candidate sites

We assigned each unit a receptive-field (RF) center at the center of its spatial position on the layer’s feature map, which spans the 8° image. For each orientation, candidate sites were units that met two conditions: (i) their RF center lay at least 2° from the grating, well outside the stimulated region; and (ii) they were genuinely driven by it (*M*_*i*_(*θ*) > 10^−4^ max_*i*,*θ*_ |*M*_*i*_(*θ*)|).

#### Iso-orientation specificity

Some units respond strongly to the central grating whatever its orientation. To isolate what each orientation in particular recruits, we ranked candidates by orientation-specific modulation, 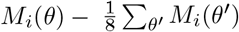. We took the top 5% as “landing sites,” identified independently for each stimulus orientation. For each landing site we computed the axial difference Δ (0–90°) between its preferred orientation and the grating orientation. We then measured the fraction of landing sites with Δ ≥ 45°. Models without landing sites were assigned “NA”. We compared these values with macaque V1, in which 30 ± 10% of labeled patches lie ≥ 45° from the injection site’s preference [15].

#### Patchiness

To quantify whether landing sites form discrete clusters on the simulated cortical sheet, we ranked candidates by raw modulation *M*_*i*_ (*θ*), took the top 5%, and computed a Clark–Evans ratio. This ratio is the mean nearest-neighbor distance between landing sites on the sheet, divided by the same quantity in null maps containing the same number of sites. We drew null sites from the candidates with probability proportional to the real sites’ density smoothed with a 1° Gaussian. Null maps therefore keep the real map’s large-scale structure but none of its clustering below that scale. A ratio of 1 indicates scattered sites and a ratio below 1 indicates patches. We averaged over 20 null maps per orientation and computed the ratio only when there were at least 10 sites.

### Model–brain mapping

#### Brain data

Neural predictivity was evaluated on the THINGS ventral spiking dataset (TVSD) [44], which contains normalized multi-unit activity recorded from two macaques (monkeys *F* and *N*) in response to images from the THINGS dataset [79]. Electrodes were localized to V1, V4, and IT based on known array implant positions, with 512 V1 channels, and 256–320 channels each in V4 and IT per monkey, concatenated across animals. The temporally averaged responses were used, yielding a single firing rate per image per electrode. The dataset came pre-split into a train and test set, with images in the test set presented 30 times to the two monkeys.

#### Mapping

Model features were mapped to trial-averaged neural responses using ridge regression with regularization strength *α* selected per electrode over *α* ∈ {10^−10^,…, 10^10^}. Predictivity was evaluated using 10-fold cross-validation and scored per electrode using the Pearson correlation. Scores were aggregated by taking the median across electrodes within each fold, then averaging across folds. An internal noise ceiling was estimated from Spearman–Brown corrected split-half consistency of the repeated neural responses.

### Lateral wiring length

For a given transformer attention layer, wiring cost was computed as the attention-weighted mean physical distance between query and key tokens. Attention matrices of shape (*B, H, Q, K*) were extracted over 1,024 images from ImageNet [36] and each attention weight *a*_*qk*_ was multiplied by the Euclidean distance *d*_*qk*_ between the corresponding token positions on the tissue map:

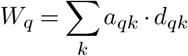

For swin transformers, distances were computed in global tissue coordinates across all attention windows, with each window’s wiring cost aggregated and averaged across windows and heads.

### Task performance

#### Image categorization performance

1000-way ImageNet categorization performance was evaluated using linear probing. Pretrained model weights were frozen and a single linear classifier was trained on top of the extracted features from the last model layer (except in the case of supervised models where features were extracted from the penultimate layer) for 30 epochs. Hyperparameters were chosen according to the evaluation configurations released by the authors of the respective model architectures. Performance was quantified as top-1 accuracy on the ImageNet validation set.

#### Action categorization performance

Kinetics-400 [50] action recognition performance was evaluated via linear probing. All pretrained weights were frozen and a linear classifier was trained for 10 epochs on top of model features extracted as mentioned above. Video clips were processed frame-by-frame, with each frame passed independently through the frozen backbone; the resulting logits were averaged across frames before computing the classification loss. Top-1 accuracy was reported on the Kinetics-400 validation set.

### Spatially-local transformers

Swin transformers [32] were trained with varying attention window sizes that determine the spatial extent over which tokens can interact. We considered three configurations, specified by the window size used in each of the four Swin transformer stages: *Large*: [7, 7, 7, 7]; *Medium*: [4, 4, 7, 7]; *Small*: [2, 2, 2, 7]. Each number denotes the side length of a square attention window, such that its square corresponds to the number of tokens within the window. Smaller window sizes therefore impose stronger locality constraints on token–token interactions, particularly in the early stages of processing. Swin transformers are trained using the self-supervised MoBY task objective [80].

### Privileged axes and privileged heads

#### Identifying privileged axes

To ask whether individual axes recur across independently trained networks, we matched units between every pair of transformer models trained with independent random seeds. Using ImageNet images, we computed the full cross-model matrix of unit response correlations on the training images and used the Hungarian algorithm to obtain a strict one-to-one assignment [81], pairing each unit in one network with a unique unit in the other. We then scored each layer by the fraction of units with a reciprocal counterpart. After fixing the assignment on the training images, an assigned pair (*j, k*) was counted only if, on held-out test images, *k* was the most correlated unit with *j* and *j* was the most correlated unit with *k*. As a chance floor, we repeated the entire procedure after independently permuting image order within each split for each network. This preserves each unit’s marginal response distribution while destroying stimulus-by-stimulus correspondence between networks.

We next characterized the visual properties encoded by these reproducible axes. We counted an axis as carrying object-property information when either shape or color explained at least 10% of its response variance, that is, when 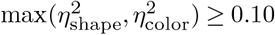. The two properties were evaluated on the same axes and are not mutually exclusive, so this measure captures the fraction of axes carrying either property rather than partitioning the axes between shape and color.

We applied a parallel criterion to orientation using a separate stimulus set of single sine gratings varying in orientation *θ*, spatial frequency *f*, phase *ϕ*, and color *χ*. Because a unit’s response to a grating depends on the phase at which the grating falls within its receptive field, raw response magnitude conflates orientation tuning with phase alignment. We therefore isolated orientation tuning using the first phase harmonic. For *P* = 4 evenly spaced phases,

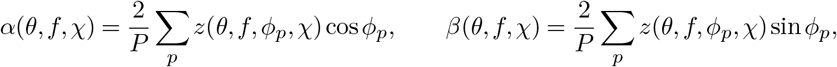

with modulation amplitude 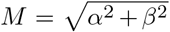, which is invariant to the grating’s absolute phase. For each unit, we identified the preferred spatial frequency and color as the pair maximizing ∑_*θ*_ *M*, and at that pair quantified orientation selectivity using

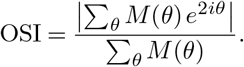

The preferred orientation was 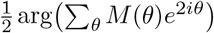 mod 180°. We counted an axis as carrying orientation information when OSI ≥ 0.5.

#### Privileged attention heads

To test whether routing is organized into reproducible components, we measured the recurrence of attention heads across independently trained transformers. For each layer and pair of models, we used the Hungarian algorithm to obtain a strict one-to-one assignment between the twelve attention heads and evaluated it using the same reciprocal held-out criterion as for units: an assigned pair passed only if each head was the other’s most similar head on the test images. We used an image-scrambled control as the reference, independently permuting image order within each split for each model.

### Readout of task-relevant representations

We assessed the linear accessibility of task-relevant information using the contour detection task [55] and the object relational reasoning task [56]. For each model and layer of interest, we extracted features from training images and validation images and flattened the resulting activations into a single feature vector for each image. We trained a logistic regression classifier using the L-BFGS solver [82]. The regularization strength *C* was selected by 5-fold cross-validation from {0.001, 0.01, 0.1, 1, 10, 100}. The classifier with the highest cross-validated accuracy was then refit on the full training set and evaluated on the held-out validation set.

### Optimal stimuli analysis

To visualize the visual features represented by transformer units, we generated feature-optimal stimuli using activation maximization following the optimization procedure of Ghiasi et al. [83]. For each analyzed layer, we randomly selected ten feature channels and optimized an input image to maximize the activation of a single feature while holding the network weights fixed. Images were optimized for 400 iterations using Adam (learning rate 0.1) with cosine learning-rate decay. To encourage invariance to nuisance image statistics and improve interpretability, each optimization step applied stochastic differentiable augmentations before evaluating the network, including random spatial jitter, color jitter, additive Gaussian noise, and repeated batch augmentation. Pixel intensities were clipped after every optimization step to remain within the valid image range.

### Statistical information

All analyses were conducted using one- or two-sided tests as specified by the directional hypothesis. Replicates were defined according to the analysis: these ranged from independent random samples of a set of images to individual models within microcircuit types to models trained with various seeds. Student’s one-sample or paired *t*-tests were used for directional comparisons and Wilcoxon signed-rank tests for paired non-parametric comparisons, while Mann–Whitney *U* tests were used for comparisons between independent model groups. Multiple comparisons were controlled using the Benjamini–Hochberg false-discovery-rate correction. Error bars represent 95% confidence intervals unless otherwise stated. Exact *n*, test statistics, and *p* values are reported in the main text next to the analyses.

## Code and data availability

Code and data to replicate the analyses described in the paper will be released publicly upon publication.

## Acknowledgements

We thank Kalanit Grill-Spector, Justin Gardner, Josh Wilson, Seda Akbiyik, Saloni Sharma, Seojin Lee, Khai Loong Aw, and members of the Stanford NeuroAI lab for helpful discussions on the manuscript.

## Supplementary Information

**Figure S1.**
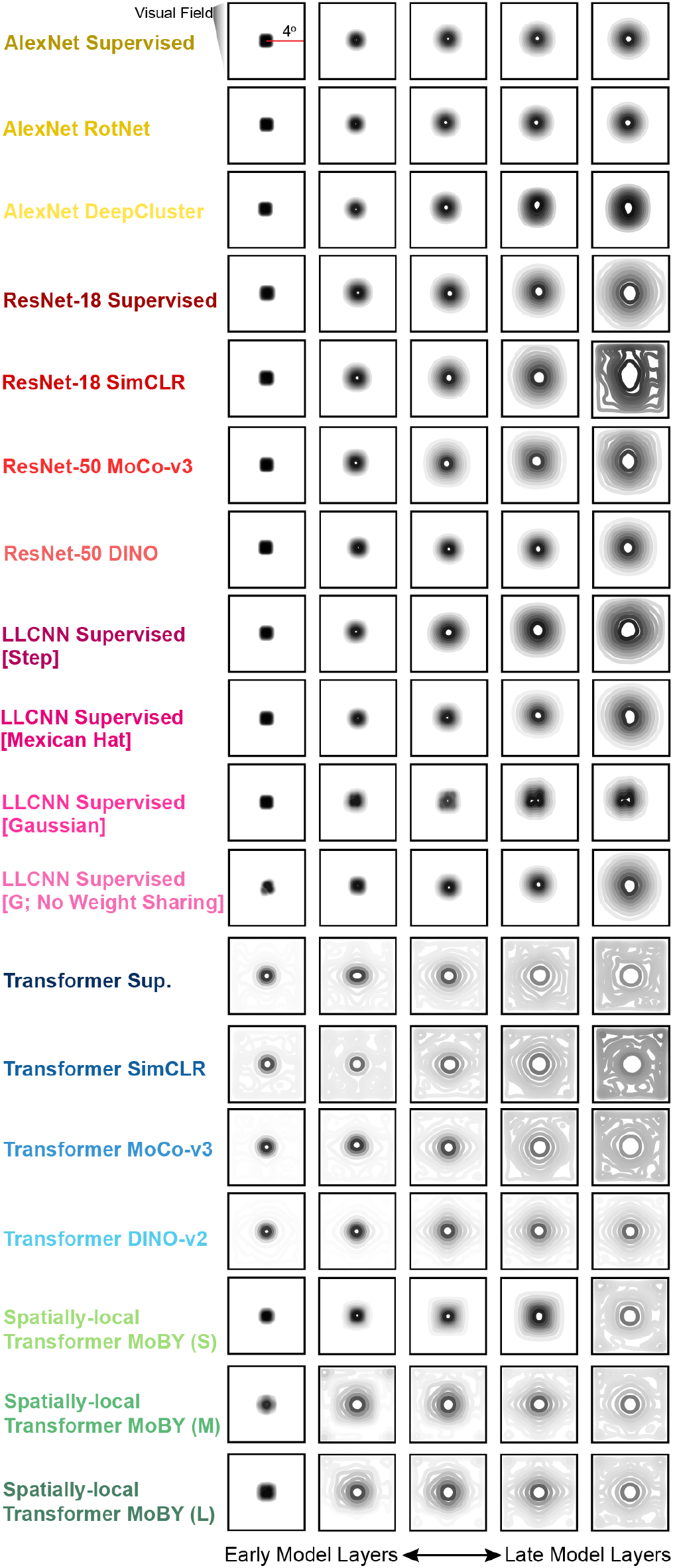
All microcircuit types exhibit an effective population receptive field hierarchy with network depth. Kernel-density estimates of effective population receptive fields (pRFs) for representative layers of each model.

**Figure S2.**
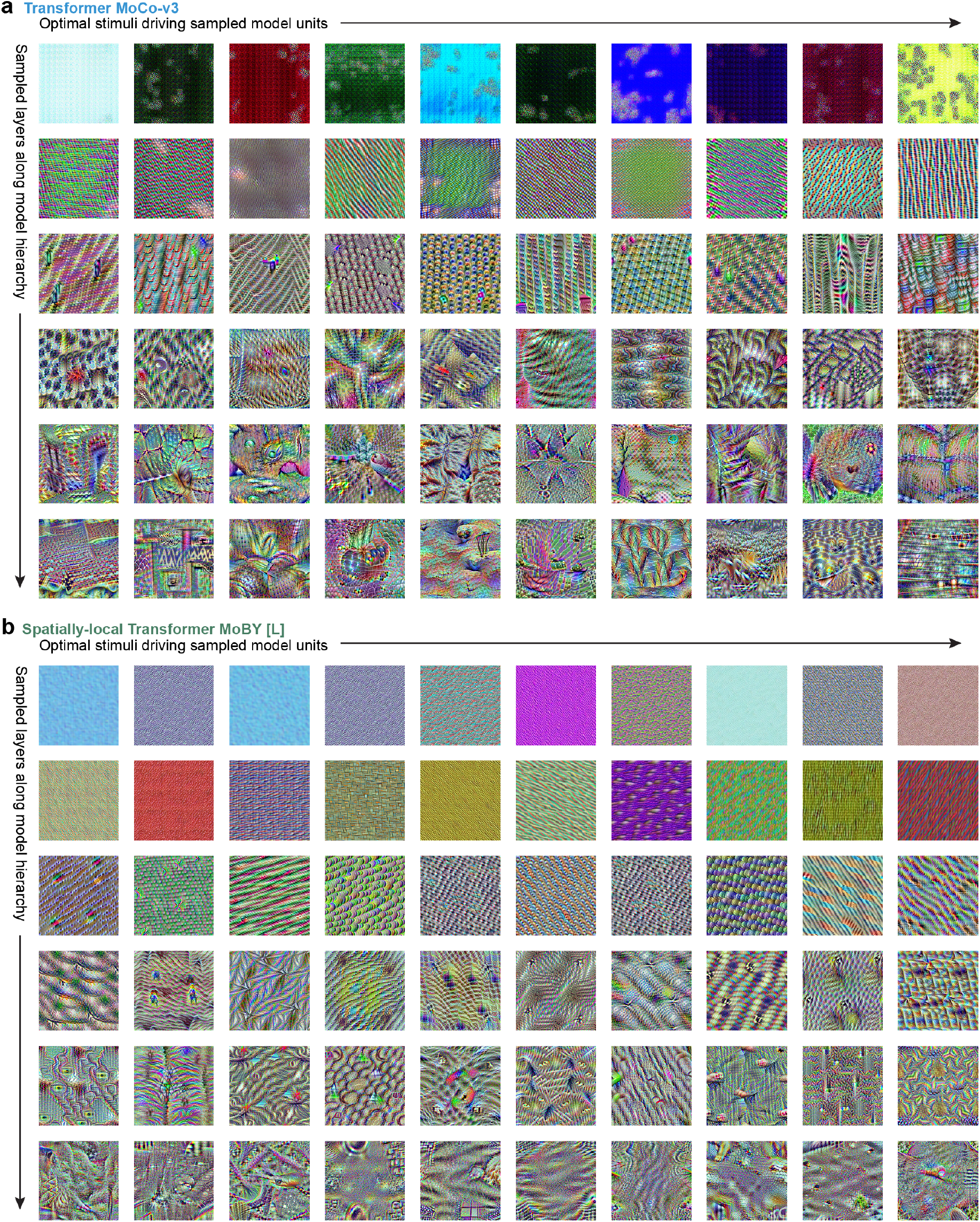
Optimal stimuli reveal hierarchical feature selectivity in transformer microcircuits, similar to convolutional networks. Representative activation-maximizing stimuli for randomly sampled units across the model hierarchy in a spatially-unconstrained transformer **(a)** and a spatially-local transformer **(b)**. For each sampled layer, ten feature dimensions were selected at random and visualized by optimizing the input image to maximize the corresponding unit’s activation while regularizing total image variation.

**Figure S3.**
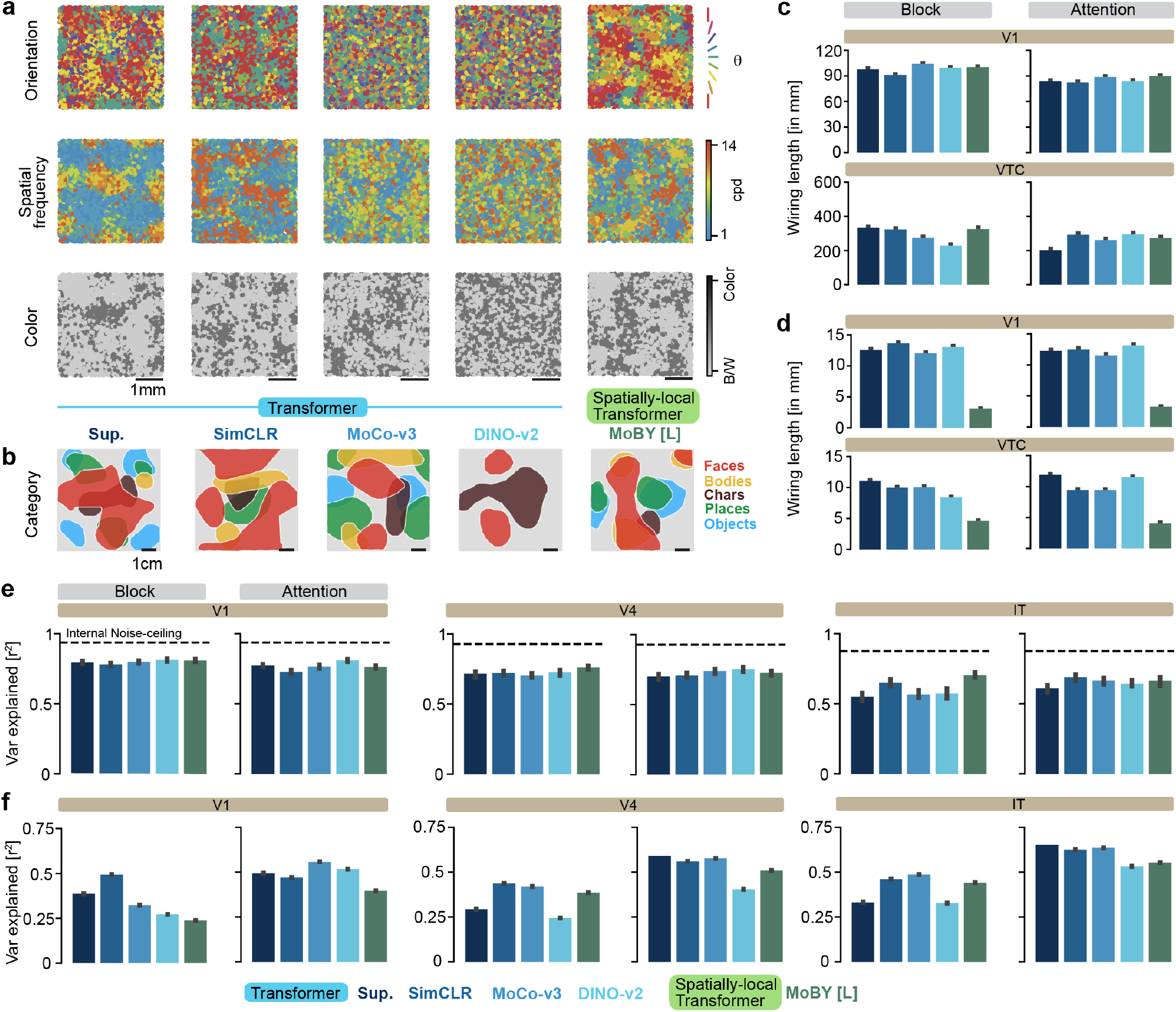
Attention outputs capture topographic and neural-response signatures rather than attention inputs. **a**. Orientation, spatial frequency, and color preference maps in the most V1-like layer of each model, using units defined as the outputs of the attention module. **b**. Same as **a**, but for smoothed category-selectivity maps in the most VTC-like layer. **c–d**. Feedforward **(c)** and lateral **(d)** wiring length in the most V1-like and VTC-like layers for alternative definitions of model units. **e–f**. Median variance explained for model-to-brain **(e)** and brain-to-model **(f)** linear mappings between the most V1-, V4-, and IT-like model layers and the TVSD dataset, using alternative definitions of model units.

**Figure S4.**
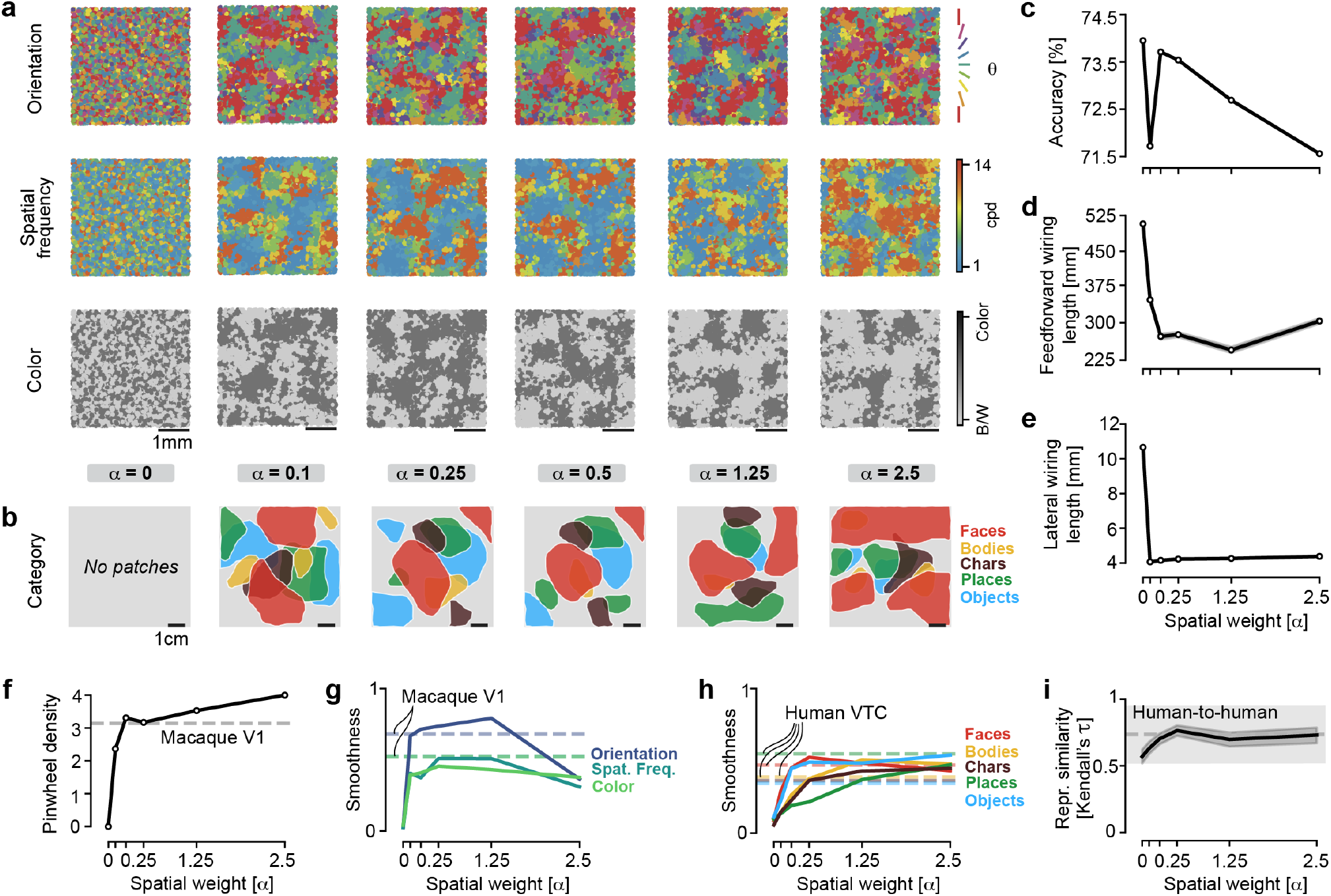
Multiple benchmarks converge on a common spatial weight for the loss function. **a**. Orientation, spatial-frequency, and color preference maps in the most V1-like layer of a Swin transformer trained with different spatial weights. **b**. Same as **a**, but for smoothed category-selectivity maps in the most VTC-like layer. **c**. Top-1 image-classification accuracy on ImageNet under a linear readout. **d–e**. Feedforward **(d)** and lateral **(e)** wiring length in the most VTC-like layer. **f**. Pinwheel density per unit area of column spacing in the most V1-like layer. *Dotted line*, macaque V1 data. **g–h**. Smoothness of V1 **(g)** and VTC **(h)** model maps. *Dotted lines*, macaque V1 and human VTC data, respectively. **i**. Functional similarity between representational similarity matrices from the most VTC-like model layer and human VTC.

**Figure S5.**
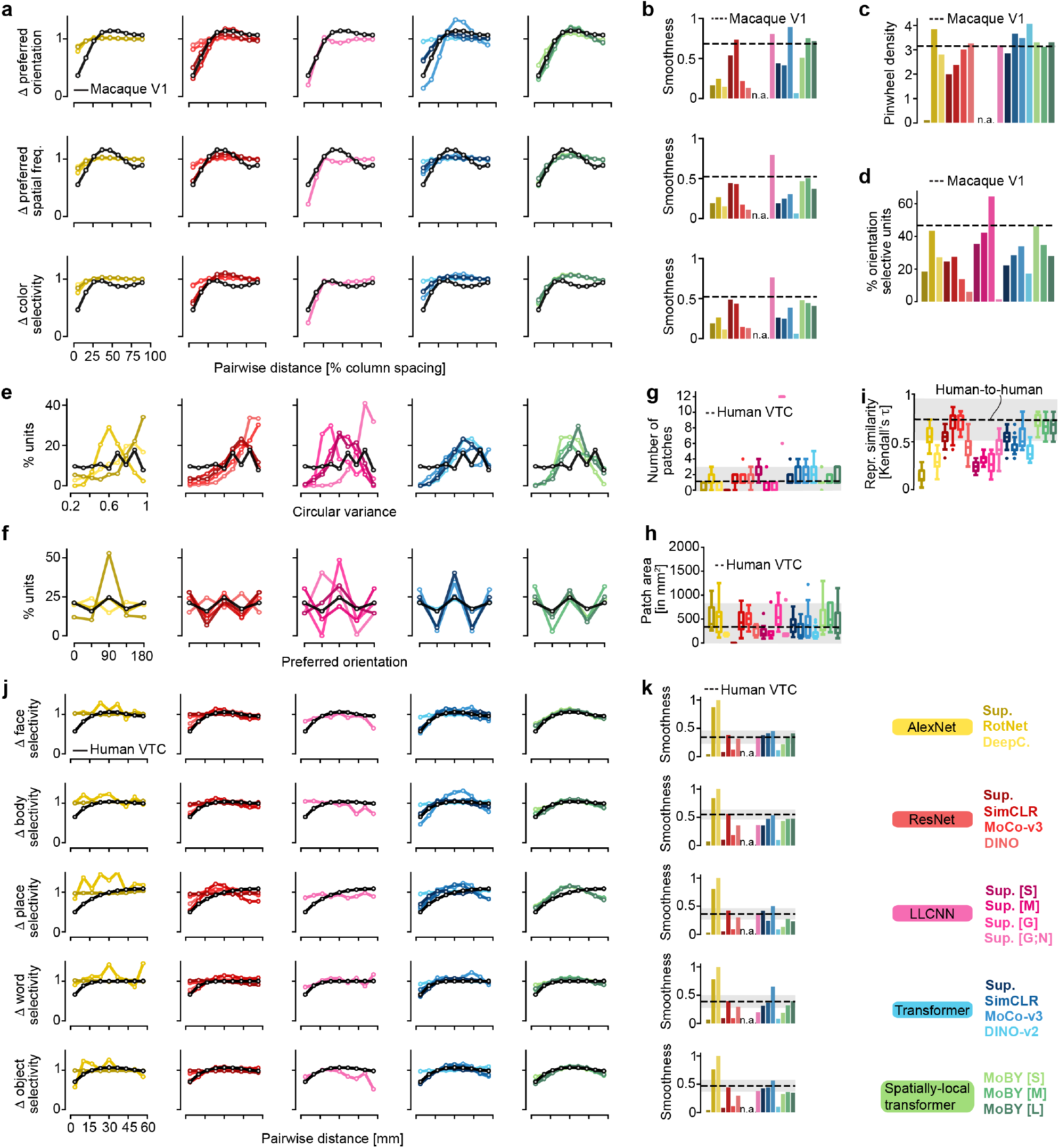
Quantitative comparison of functional tuning and organization across models. **a**. Change in preferred orientation, spatial frequency, and color as a function of cortical distance, normalized to chance, in the most V1-like layer of each model. *Black*, macaque V1 data. **b**. Map smoothness for orientation, spatial frequency, and color. **c**. Pinwheel density per unit area of column spacing in the most V1-like layer of each model. **d**. Fraction of orientation-selective units in the most V1-like layer. *Black*, macaque V1 data. **e**. Distribution of circular variance across units in the most V1-like layer. **f**. Distribution of preferred orientations across units in the most V1-like layer. **g**. Mean number of patches in VTC topographic maps. *Black*, human VTC data. **h**. Mean area covered by patches in VTC topographic maps. **i**. Functional similarity between representational similarity matrices from each model’s most VTC-like layer and human VTC. *Black*, human-to-human similarity. **j**. Change in category selectivity as a function of cortical distance, normalized to chance, for pairs of units in the most VTC-like layer. **k**. Map smoothness for category selectivity.

**Figure S6.**
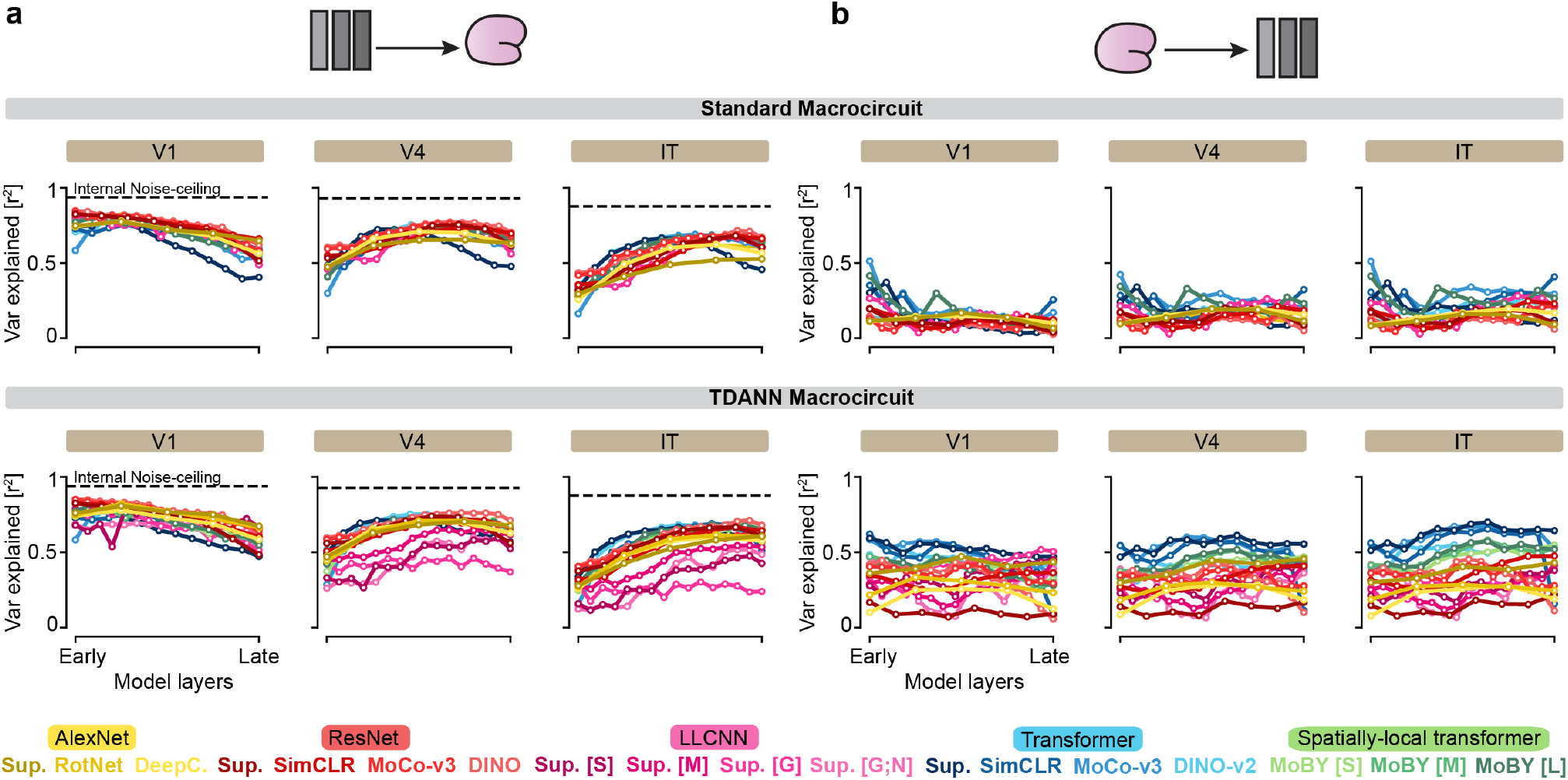
Models trained within the TDANN macrocircuit show substantially higher neural predictivity than those trained under the standard macrocircuit for brain-to-model alignment. **a**. Median variance explained by a linear mapping from model layers to TVSD responses. *Black line*, Spearman–Brown-corrected split-half consistency. **b**. Same as **a**, but for the reverse mapping from brain responses to model representations.

**Figure S7.**
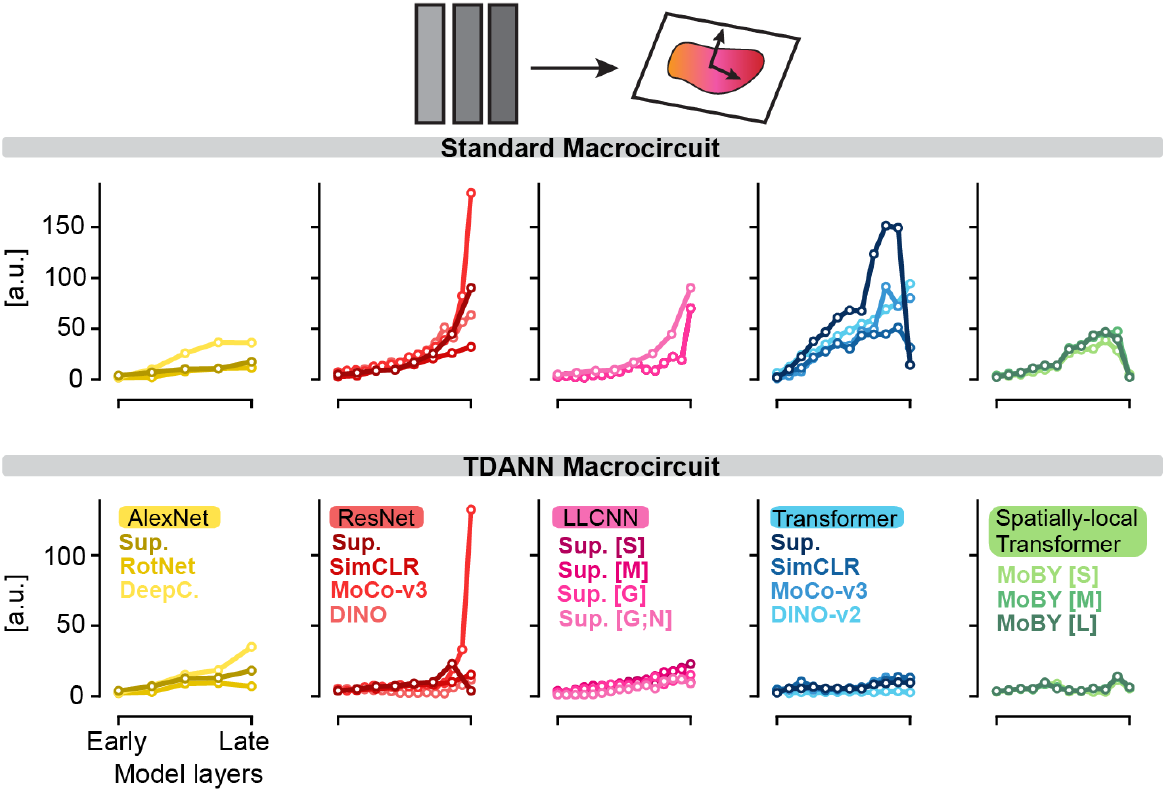
Intrinsic dimensionality across model layers. Intrinsic dimensionality computed from model representations of 1,024 images randomly sampled from the training images presented to the two monkeys in the TVSD dataset.

**Figure S8.**
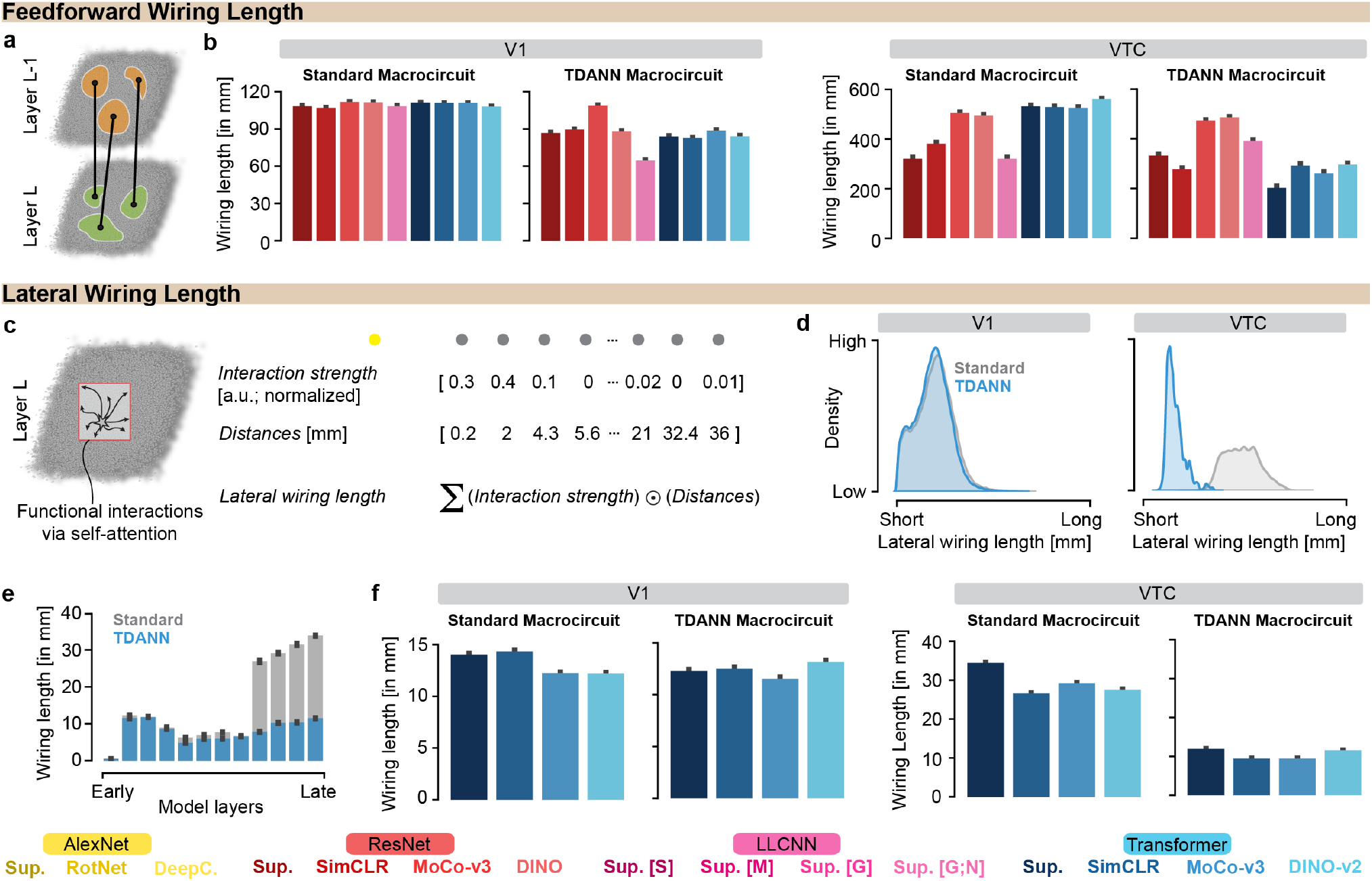
Transformers exhibit reduced feedforward and lateral wiring length. **a**. Schematic of feedforward, or between-layer, wiring. **b**. Feedforward wiring length in the most V1-like and VTC-like layers of each model, computed functionally over 1,024 randomly sampled ImageNet images. *Error bars*, 95% CI across cortical neighborhoods and input images. AlexNet and weight-sharing LLCNNs are omitted because of insufficient depth or coarse retinotopy. **c**. Schematic of lateral, or within-layer, wiring. **d**. Distribution of lateral wiring lengths for units in representative V1-like and VTC-like layers. **e**. Lateral wiring length across layers of a representative model. *Error bars*, 95% CI across model units. **f**. Lateral wiring length in the most V1-like and VTC-like layers of each model. *Error bars*, 95% CI across model units. LLCNNs are omitted because their lateral wiring length is fixed during training.

**Figure S9.**
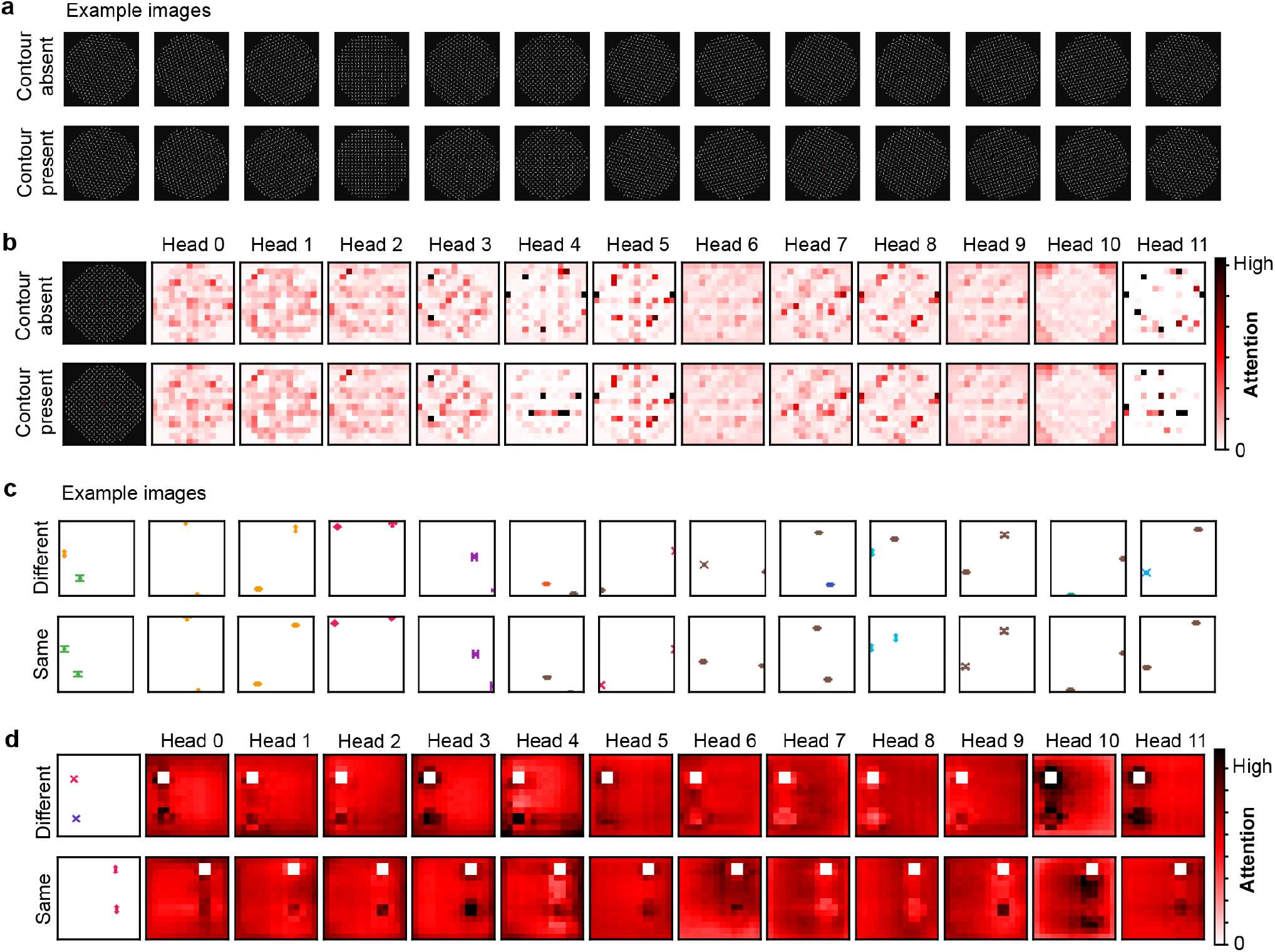
Visualizing all transformer heads and their routing patterns for both contour detection and object relational reasoning tasks. **a**. Example stimuli from the contour-detection task. **b**. Transformer-head routing from one segment of the contour to the remainder of the image. **c**. Example stimuli from the object relational reasoning task. **d**. Transformer-head routing from one object to the remainder of the image.

